# Riboswitch Folds to *Holo*-Form Like Structure Even in the Absence of Cognate Ligand at High Mg^2+^ Concentration

**DOI:** 10.1101/2021.10.05.463230

**Authors:** Sunil Kumar, Govardhan Reddy

## Abstract

Riboswitches are non-coding RNA that regulate gene expression by folding into specific three-dimensional structures (*holo*-form) upon binding by their cognate ligand in the presence of Mg^2+^. Riboswitch functioning is also hypothesized to be under kinetic control requiring large cognate ligand concentrations. We ask the question under thermodynamic conditions, can the riboswitches populate *holo*-form like structures in the absence of their cognate ligands only in the presence of Mg^2+^. We addressed this question using thiamine pyrophosphate (TPP) riboswitch as a model system and computer simulations using a coarse-grained model for RNA. The folding free energy surface (FES) shows that with the initial increase in Mg^2+^ concentration ([Mg^2+^]), TPP AD undergoes a barrierless collapse in its dimensions. On further increase in [Mg^2+^], intermediates separated by barriers appear on the FES, and one of the intermediates has a TPP ligand-binding competent structure. We show that site-specific binding of the Mg^2+^ aids in the formation of tertiary contacts. For [Mg^2+^] greater than physiological concentration, AD folds into its *holo*-form like structure even in the absence of the TPP ligand. The folding kinetics shows that it populates an intermediate due to the misalignment of the two arms in the TPP AD, which acts as a kinetic trap leading to larger folding timescales. The predictions of the intermediate structures from the simulations are amenable for experimental verification.

## Introduction

The mechanism of gene regulation by riboswitches in bacteria is a fascinating problem. Riboswitches are non-coding RNA present in the 5′ untranslated region of the mRNA and are composed of two domains: the aptamer domain (AD) and the expression platform (EP). An interplay of the structural transitions between the AD and EP play a critical role in gene regulation. ^1–3^ Structural transitions between the two domains are influenced by Mg^2+^ and the binding of a cognate ligand to the AD.

The Mg^2+^ influence RNA folding through diffuse and site-specific binding.^4^ In diffuse binding, Mg^2+^ non-specifically binds to the negatively charged phosphate groups and renormalizes charge on the RNA chain^5^ leading to the initial collapse. Whereas in site-specific binding, Mg^2+^ bind in the pockets created by the specific arrangement of nucleotides and aid in the stabilization of secondary and tertiary structures.^6,7^ However, for the functioning of riboswitches, in addition to the Mg^2+^, cognate ligand binding is equally essential. The cognate ligand binding to the AD leads to structural changes in both the AD and EP domains, which signal the attenuation or initiation of the transcription or translation process. Riboswitches are important drug targets since they are involved in the critical functional role of gene regulation in bacteria and also absent in the human genome.^8,9^

Experiments^10–22^ and simulations^23–31^ probing the folding of ADs in the presence of Mg^2+^ and their cognate ligands provide evidence that the ADs populate multiple intermediates in their folding pathways. Functional studies on TPP,^32^ Adenine,^33^ Mg^2+^,^34^ c-di-GMP,^35^ SAM-I^36^ and FMN^37,38^ riboswitches further demonstrate that higher concentrations of cognate ligands are required than their intrinsic affinity to effectively regulate gene expression, indicating that riboswitch functioning is under kinetic control rather than thermodynamic control. Despite significant progress in our understanding of the functioning of riboswitches, the role of cations and cognate ligands in the mechanism of structural transitions in the AD and EP is not completely clear. An interesting question that requires comprehensive understanding is if Mg^2+^ alone is sufficient to populate the *holo*-form like structures in the absence of the cognate ligand under thermodynamic conditions? To address this question, we studied the role of [Mg^2+^] on the stability of intermediates populated in the folding free energy surface (FES) of the AD of thiamine pyrophosphate (TPP) sensing riboswitch.

The TPP riboswitch is widely distributed along the three phylogenic branches: bacteria, archaea, and eukarya. It is found in 48 out of the 59 human bacterial pathogens making it the most abundant riboswitch in human pathogens.^39^ The TPP AD is *≈* 80 nt long, and in the native state (*holo*-form), it is a junction-type riboswitch with a tuning-fork-like structure^40^ composed of two arms (Figure 1A and S1 in supporting information (SI)). The P_2*−*3_ arm is composed of P_2_-P_3_ helices joined by the junction J_32_, and the P_4*−*5_ arm is composed of P_4_-P_5_ helices joined by the junction J_45_, respectively. The P_1_ helix acts as a base holding both the arms from P_2_ and P_4_ helices (Figure 1A). The P_2*−*3_ arm has the binding pocket for the 4-amino-5-hydroxymethyl-2-methyl-pyrimidine (HMP) moiety of the TPP ligand. The P_4*−*5_ arm binds to the pyrophosphate (PP) moiety of the ligand through the mediation of Mg^2+^.

**Figure 1:**
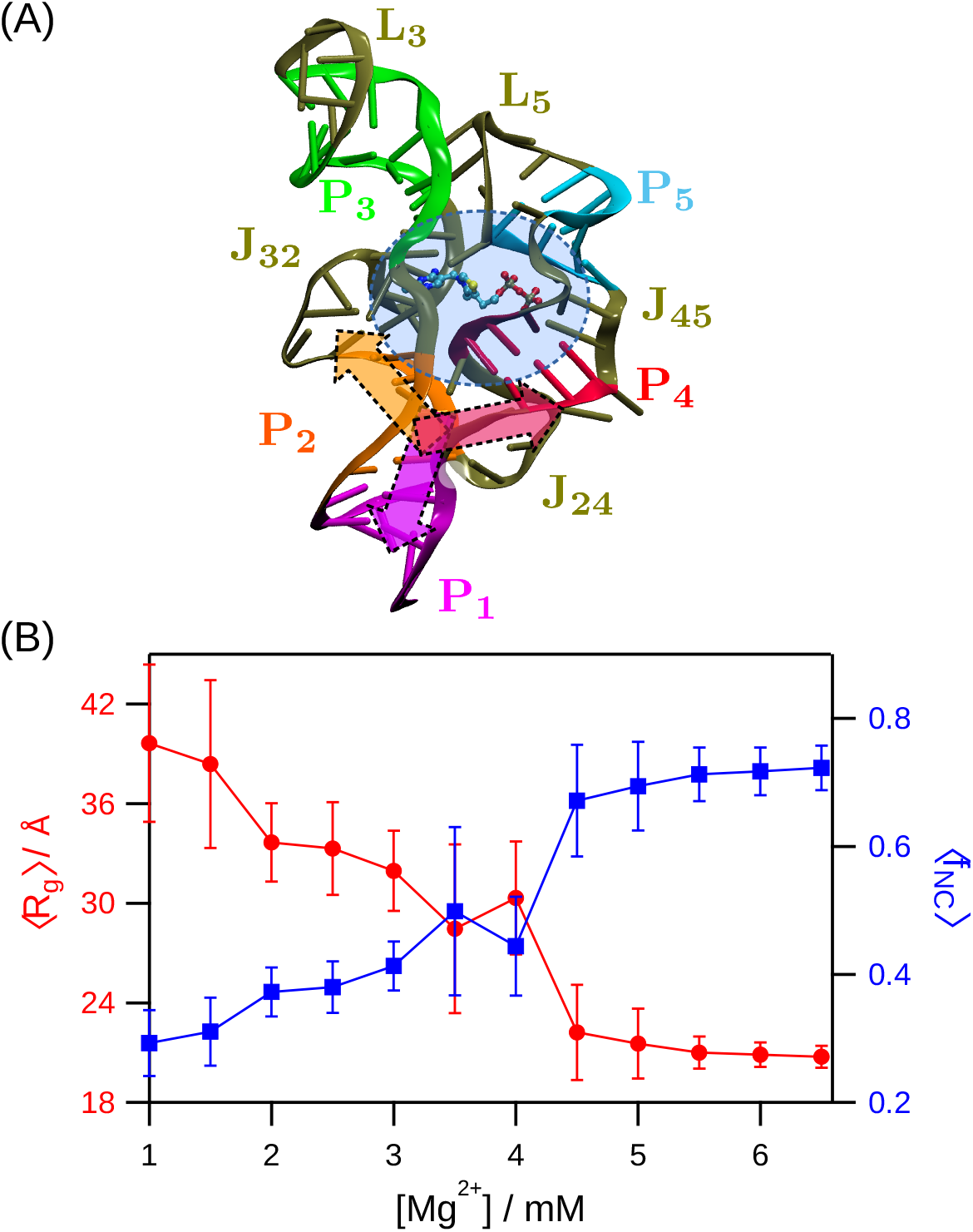
(A) Structure of the TPP AD (PDB: 2GDI^40^). Helices P_1_ to P_5_ are shown in magenta, orange, green, red, and cyan, respectively. Helical junctions (J_32_, J_45_ and J_24_) and loops (L_3_ and L_5_) are shown in tan. The core region of the AD containing the TPP ligand binding pocket is shown as a shaded area in blue. The bound TPP-ligand inside the core region is shown using ball-stick representation. The three-way junction (3WJ) is shown by the Y-shaped shaded area near the joint of P_1_ and P_2_ helices. (B) The average radius of gyration, ⟨*R*_*g*_⟩ and the average fraction of native contacts, ⟨*f*_*NC*_⟩ are plotted as a function of [Mg^2+^] using solid red circles and blue squares, respectively. Multiple transitions in both of the plots indicate that TPP AD is a multi-state folding system.

Single-molecule FRET^12,18^ and SAXS^41–43^ studies have shown that physiological [Mg^2+^] is indispensable for the cognate ligand to bind to the TPP AD. Single-molecule FRET^18^ study proposed that a Y-shaped intermediate mediated by Mg^2+^ gets populated during the initial stages of folding even in the absence of the cognate TPP ligand. Upon binding of the TPP ligand, the AD attains the prerequisite folded state that regulates the downstream gene expression. Studies on other riboswitches reveal that the AD domains can sample the *holo*-form like conformations even in the absence of cognate ligand but in [Mg^2+^] greater than the physiological concentration.^44,45^ Experiments studying TPP AD folding also suggest that multiple structural ensembles are populated depending on [Mg^2+^] and TPP-ligand.^12^ Further, the timescales associated with the transitions among these ensembles varied from *µ*s to s, indicating significant variations in the barrier heights separating the basins.

Computer simulations using all-atom^7,30,46–51^ and coarse-grained models^52–61^ are playing an important role in elucidating various aspects of RNA folding. The stability of folded state compared to the unfolded state of RNA is very sensitive to the charge and the concentration of the cations.^62,63^ Mg^2+^ is known to be remarkably efficient in facilitating the formation of native-like tertiary contacts and the folded state even in the millimolar concentration. In contrast, submolar to molar concentration of K^+^ is required for the monovalent ion-driven folding of RNA.^63–68^ Mg^2+^ preferentially accumulates around the phosphate groups (P sites) of specific nucleotides and initiates folding by stabilization of the secondary structures and subsequently leading to the hierarchical formation of tertiary contacts.^56,69,70^ During various stages of RNA folding, the inherently hierarchical nature of structural reorganization can result in the population of intermediates with an increase in [Mg^2+^].

We studied the effect of [Mg^2+^] on the folding FES of TPP AD using molecular dynamics simulations and a coarse-grained RNA model.^56^ We find that with the increase in [Mg^2+^], AD initially undergoes barrierless compaction in size followed by the formation of intermediate states. We show that Mg^2+^ binding to the P sites of specific nucleotides leads to the formation of tertiary contacts that stabilize the *holo*-form like folded state even in the absence of the TPP ligand. Interestingly, folding kinetics show that the AD can populate a kinetic trap due to the misaligned orientation of the arms leading to a larger folding time.

## Methods

We studied the effect of [Mg^2+^] on the folding mechanism of TPP AD from *Escherichia coli thiM* mRNA using the three interaction site (TIS) RNA model^56,71^ and Langevin dynamics simulations. The TIS model for the TPP AD is constructed using the crystal structure (PDB: 2GDI).^40^

### TIS Model of RNA

We used the TIS model developed by Denesyuk and Thirumalai,^56,71^ where each nucleotide is represented by three sites mimicking the phosphate (P), sugar (S), and base (B) groups. The centers of the sites are placed at the center of mass of the P, S, and B groups, respectively. In this model, the monovalent (K^+^, Na^+^, and Cl^*−*^) and divalent ions (Mg^2+^) are explicitly present in the simulation. The Hamiltonian for the TIS model has seven components and is given by

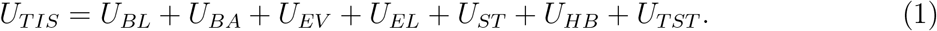

The components in the Hamiltonian correspond to the bond length (*U*_*BL*_), bond angle (*U*_*BA*_), excluded volume repulsion between different sites (*U*_*EV*_), electrostatic interaction between charged sites (*U*_*EL*_), single-strand base stacking interaction between consecutive bases (*U*_*ST*_), hydrogen bonding interaction (*U*_*HB*_) and tertiary stacking interaction between two non-consecutive bases (*U*_*TST*_), respectively. The force field details and parameters can be found in the works of Denesyuk and Thirumalai.^56,71^ Below we provide a brief description of the force field.

The harmonic bond length and bond angle potentials account for the RNA chain connectivity and stiffness. The excluded volume repulsion between a pair of interacting sites, either RNA or ions, is given by a modified Lennard-Jones potential. The electrostatic interaction between a pair of charged sites is given by the Coulomb potential scaled by the temperature-dependent dielectric constant of water. The charge is -1 for phosphate group, -1 for Cl^*−*^, +2 for Mg^2+^, +1 for K^+^, and +1 for Na^+^ ions, respectively. The consecutive bases in the RNA chain have base stacking interactions, which depend on the sequence. The list of base pairs involved in the native hydrogen bond network in the folded structure of TPP AD is obtained using the crystal structure (PDB: 2GDI) and WHAT IF web server (https://swift.cmbi.umcn.nl) ^72^(Table S1-S4). All the base pairs, which have a hydrogen bond between them, interact using the hydrogen bonding potential. Non-canonical base pair hydrogen bonds and tertiary base stacking interactions present in the crystal structure are also taken into account in the model (Table S5-S6). This model is successful in quantitatively accounting for the folding thermodynamics of various RNA systems such as *Azoarcus* ribozyme,^56^ the central domain of 16S ribosomal RNA^70^ and RNA pseudoknots^68^ demonstrating that it is reliable and transferable to study various other RNA systems.

### Simulations

We performed simulations to study TPP AD folding as [Mg^2+^] is varied from 1 mM to 6.5 mM. The [K^+^] is fixed at 30 mM in all the simulations. The simulations are performed in a cubic box of length 200 Å. The two Na^+^ ions present in the crystal structure are added to the simulation box. The number of Mg^2+^ and K^+^ ions in the simulation box are computed using their concentration and box volume. Cl^*−*^ ions are added to maintain charge neutrality in the simulation box. All ions are explicitly modeled as beads with charge and excluded volume. We used Langevin dynamics simulations to study the folding dynamics of TPP AD at temperature, *T* = 310 K. To compute the thermodynamic properties of TPP AD folding, we used 5% viscosity of water, *η* = 5 *×* 10^*−*5^ Pa*·*s to enhance conformational sampling. To study the TPP folding kinetics at [Mg^2+^] = 6.5 mM, we used the viscosity of water, *η* = 10^*−*3^ Pa*·*s. The equation of motion for the RNA sites and ions in Langevin dynamics is given by

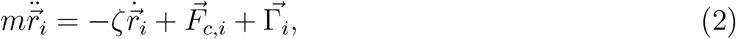

where *m*_*i*_ is the mass (in amu), *ζ*_*i*_ (= 6*πηR*_*i*_) (in amu/fs) is the friction coefficient, *r*_*i*_ is the position, 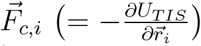 is the deterministic force, *R*_*i*_ is the radius (in Å) of *i*^*th*^ site in the system. 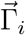 is the random force on the *i*^*th*^ site with a white-noise spectrum. The random force auto-correlation function is given by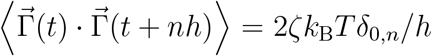, where *n* = 0, 1, …, *δ*_0,*n*_ is Kronecker delta function, and *k*_B_ is the Boltzmann constant. The Langevin equation is integrated using the velocity Verlet algorithm with a time step *h* (= 2.5 fs).^56,73^ System coordinates are saved after every 5,000 steps (*τ*_*f*_ = 12.5 ps) to compute the properties. The initial 1 *µ*s of simulation data is ignored in computing the properties. For each [Mg^2+^], at least ≈ 13 *µ*s of simulation data is collected to compute the average properties. The atomistic coordinates of the AD are generated using the coarse-grained coordinates and TIS2AA^74^ program, which uses the fragment-assembly approach^75^ and energy minimization embedded in AmberTools.^76^ We used VMD to generate the three dimensional structures of the AD.^77^

### Data Analysis

The radius of gyration *R*_*g*_, of the TPP AD is computed using the equation, 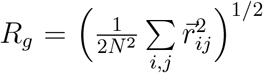, where *N* (= 240) is the total number of RNA sites in the TIS model of TPP AD, and 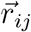 is the vector connecting the sites *i* and *j*. The average fraction of helix formation for a given helix *H*, is computed using the equation

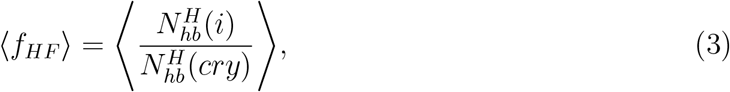

where 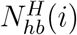 is the number of hydrogen bonds present in the helix *H* in *i*^*th*^ conformation and 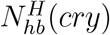 ⟨ ⟩is the total number of hydrogen bonds present in the helix *H* in the crystal structure. denotes the average over all the conformations. A hydrogen bond is considered to be present if its energy is lower than the thermal energy (*k*_B_*T*).^56^

### Local [Mg^2+^] Around P sites

The local [Mg^2+^] in the vicinity of the P site of the *i*^*th*^ nucleotide in molar units is computed^69^ using the relation

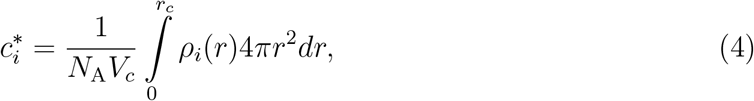

where *ρ*_*i*_(*r*) is the number density of the Mg^2+^ at a distance *r* from the *i*^*th*^ P site, *V*_*c*_ is the spherical volume of radius *r*_*c*_, and *N*_A_ is the Avogadro’s number. The cutoff radius *r*_*c*_ is given by *r*_*c*_ = *R*_*Mg*_ + *R*_*P*_ + Δ*r*, where *R*_*Mg*_ and *R*_*P*_ are the radii of Mg^2+^ and P sites, and Δ*r* is the margin distance. We used Δ*r* = 1.7 Å to ensure that we take into account only the tightly bound local Mg^2+^ around the P sites.

### FES Calculation

The total number of native contacts in the TPP AD is computed using the TIS model of the folded crystal structure (PDB: 2GDI).^40^ A pair of sites *i* and *j* in the folded structure are defined to have a native contact between them if |*i − j*| *>* 10, and the distance between the sites *r*_*ij*_ is less than 15 Å (Figure S1). The *f*_*NC*_ for *i*^*th*^ TPP AD conformation is computed using the equation

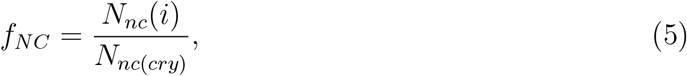

where *N*_*nc*_(*i*) the number of native contacts present in *i*^*th*^ conformation and *N*_*nc*_(*cry*) is the number of native contacts present in the crystal structure.

The FES corresponding to the TPP AD folding (*G*) is projected onto the fraction of native contacts *f*_*NC*_, and it is calculated using the equation

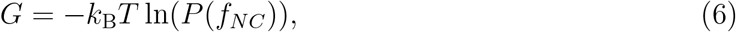

where *P* (*f*_*NC*_) is the probability distribution of *f*_*NC*_.

### Fraction of Tertiary Contacts (TCs)

The TPP AD has two critical tertiary contact (TC) forming sites, the three-way junction (TC_3*WJ*_) and the arm-tip (TC_*AT*_). The TC_3*WJ*_ involves contacts between the nucleotide segments A12 - U20, C48 - U59, and A80 - G86 (Figure S2A). The TC_*AT*_ involves contacts between the nucleotide segments G21 - C24, and A69 - G72 (Figure S2A). Apart from the two TCs mentioned above, TPP AD has a third tertiary contact (TC_*RB*_) between U54 (J_24_) and U79 (J_45_) located at the base of the P_4*−*5_ arm. The total number of tertiary interactions in a particular TC in the TPP AD native state is equal to the number of native contacts between the sites belonging to the nucleotides involved in forming that specific TC. The fraction of tertiary contacts *f*_*TC*_ for a particular TC is computed using the equation

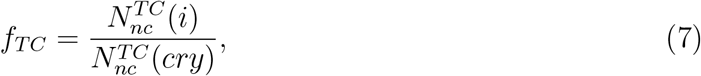

where 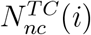 is the number of native contacts present between the sites belonging to the nucleotides involved in the formation of that specific TC in the *i*^*th*^ conformation and 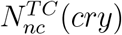 is the total number of native contacts present in that specific TC in the crystal structure. We label a TC as formed (F) if ⟨*f*_*TC*_ ⟩ is *≥* 0.5 and as ruptured (R) otherwise.

### Role of Mg^2+^ in Tertiary Contact Formation

The contribution of Mg^2+^ binding to the P sites in the formation of TCs is inferred by computing the free energy difference (ΔΔ*G*_*TC*_) in the TC formation with and without Mg^2+^ binding.^70^ The free energy contribution to the TC formation due to Mg^2+^ binding to the *i*^*th*^ P site is given by

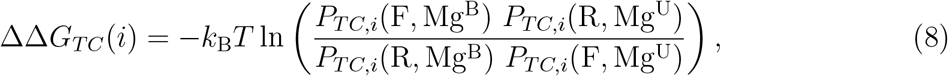

where *P*_*TC,i*_(F, Mg^B^) is the joint probability that TC is formed (F), and Mg^2+^ is bound (Mg^B^) to the *i*^*th*^ P site, *P*_*TC,i*_(R, Mg^U^) is the joint probability that TC is ruptured (R) and Mg^2+^ is not bound (Mg^U^) to the *i*^*th*^ P site. Similarly *P*_*TC,i*_(R, Mg^B^) and *P*_*TC,i*_(F, Mg^U^) are defined. An Mg^2+^ is considered bound to the *i*^*th*^ P site if the distance between the centers of the P site and ion is less than 4.6 Å.

### TPP AD Folding Kinetics

To decipher the kinetic intermediates populated in the TPP AD folding pathways, we computed an inter-arm angle (Ω_*IA*_) between the helical arms to characterize their orientation. The arm with helices P_2_ and P_3_ is represented using a vector 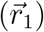 joining the center of mass of nucleotides C15 and G51, and the center of mass of nucleotides C22 and G37. Similarly, the other arm with helices P_4_ and P_5_ is represented using a vector 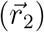 joining the center of mass of nucleotides C57 and G82, and the center of mass of nucleotides U64 and A75. The inter-arm angle Ω_*IA*_ is defined to be the angle between the vectors 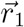 and 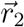(Figure S2B). We computed a dihedral angle (Θ_*J*_) to characterize the orientation of the junction J_24_. Θ_*J*_ is computed using the position of the S sites of the nucleotides U14, G51, C57 and G86 belonging to P_1_, P_2_, P_4_, and P_1_ helices, respectively. The length of J_24_ motif 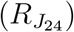 is the distance between the P sites of the nucleotides G51 and C57 (Figure S2B).

## Results

### Mg^2+^ Facilitates TPP AD Folding

In physiological conditions, noncoding RNA molecules fold to a unique native state to perform their regulatory activities. RNA molecules have a cooperative interaction network,^78,79^ which guides folding to the native state and suppresses the formation of non-native structures from the early stages of folding.^80^ The cooperativity in RNA folding further depends on the ions present in solution.^81–89^ To study the effect of Mg^2+^ on TPP AD folding, we computed ⟨*R*_*g*_ ⟩ and ⟨*f*_*NC*_⟩ as a function of [Mg^2+^] (Figure 1B). As we increase the [Mg^2+^] from 1 mM to 6.5 mM, the TPP AD undergoes a transition from an unfolded state (*R*_*g*_ ≈ 40 Å and *f*_*NC*_ ≈ 0.3) to a compact folded state (⟨*R*_*g*_ ⟩ ≈ 21 Å and ⟨ *f*_*NC*_ *≈ ⟩* 0.7). The plot shows two major folding transitions (Figure 1B). The first transition occurs in the range 1.5 mM < [Mg^2+^] < 2 mM, where ⟨ *R*_*g*_ ⟩ decreases from ≈ 38 Å to 34 Å (*f*_*NC*_ changes from 0.31 to 0.37), and the second transition occurs in the range 4 mM < [Mg^2+^] < 4.5 mM, where ⟨*R*_*g*_⟩ decreases from ≈ 30 Å to 22 Å (⟨*f*_*NC*_⟩ changes from 0.44 to 0.67). The multiple transitions indicate that TPP AD populates intermediates when folding is initiated by increasing [Mg^2+^]. The changes in ⟨*R*_*g*_⟩ and ⟨*f*_*NC*_⟩ with the increase in [Mg^2+^] shows that TPP AD can fold to its native-like state only beyond physiological [Mg^2+^] (≈ 4 mM^90–92^) in the absence of the TPP ligand.

### Folding Intermediates and *Holo*-Form Like Structure

The FES projected onto *f*_*NC*_ shows that AD populates four different states depending on the [Mg^2+^] (Figure 2). As [Mg^2+^] is increased, the equilibrium shifted towards populating compact intermediate states. Experiments have shown that Mg^2+^ contributes to both specific and non-specific compaction in the dimensions of the RNA unfolded state.^18,41,85,93,94^ In the early stages of folding, we observed barrierless collapse of the AD as [Mg^2+^] is increased from 1 mM to 2 mM, where the FES retains its shape and only its minimum is shifted from *f*_*NC*_ = 0.29 to 0.38, indicating compaction in the AD size (Figure 2). The decrease in ⟨*R*_*g*_ ⟩ and the disrupted tertiary contacts confirm compaction in the AD size without the formation of any tertiary structure (Figure 1B and 3A). For [Mg^2+^] *≤* 2 mM, the AD unfolded state is the global minimum on the FES. Upon increasing [Mg^2+^] (*≥* 3 mM), intermediate states separated by energy barriers appear on the FES along with the shift in energy minima to higher *f*_*NC*_. For [Mg^2+^] *≥* 5 mM, the global minimum on the FES is the AD native-like folded state (*holo*-form) with *f*_*NC*_ *≥* 0.7.

**Figure 2:**
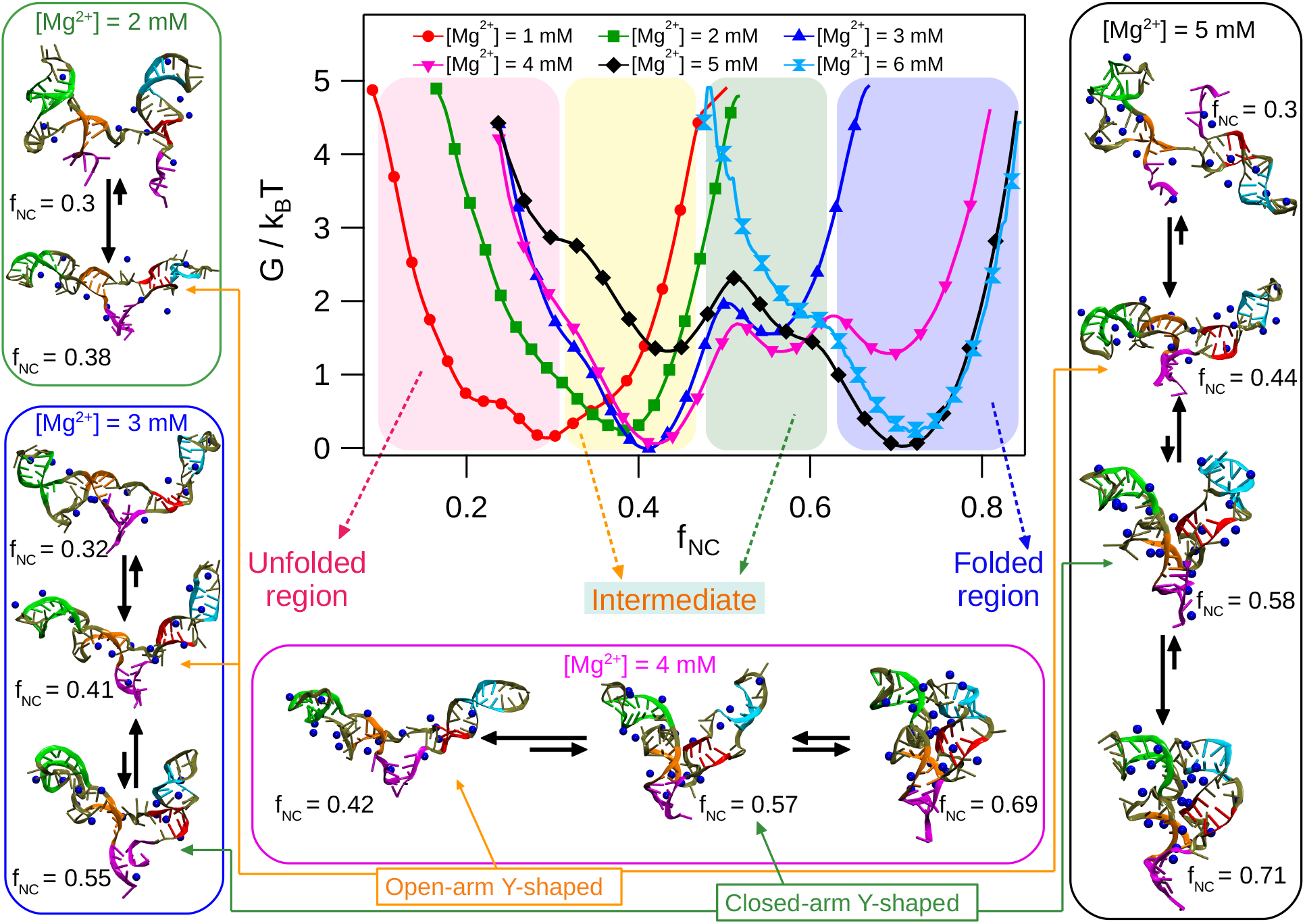
FES of the TPP AD projected onto the fraction of native contacts, *f*_*NC*_. The FES for [Mg^2+^] = 1 to 6 mM are plotted using solid red circle, green square, blue triangle, magenta invertedtriangle, black rhombus, and cyan hour-glass markers, respectively. The FES is divided into four regions. The unfolded and folded regions are shown in red and blue. The region showing intermediates with open and closed-arm Y-shaped structures are shown in yellow and green. Representative conformations of the AD in each basin of the FES are shown in cartoon representation. P_1_ to P_5_ helices are shown in magenta, orange, green, red, and cyan colors, respectively. The condensed Mg^2+^ on the TPP AD are shown as blue beads. The equilibrium between structures in different basins are shown for different [Mg^2+^].

For [Mg^2+^] = 3 mM, the TPP AD exists in three states: unfolded (*f*_*NC*_ = 0.32), open-arm Y-shaped (*f*_*NC*_ = 0.41) and closed-arm Y-shaped (*f*_*NC*_ = 0.55) (Figure 2). As [Mg^2+^] increases to 5 mM, the FES shows that the TPP AD populates four states: unfolded (*f*_*NC*_ = 0.3), open-arm (*f*_*NC*_ = 0.44), closed-arm (*f*_*NC*_ = 0.58) and *holo*-form like folded state (*f*_*NC*_ = 0.71) (Figure 2). However, with the increase in [Mg^2+^] from 3 mM to 5 mM, the most stable state shifts from the open-arm Y-shaped (*f*_*NC*_ = 0.41) to the *holo*-form state (*f*_*NC*_ = 0.71) (Figure 2). The increase in stability of the *holo*-form like state (*f*_*NC*_ = 0.71) for [Mg^2+^] *≥* 4.5 mM is due to the formation of both TC_3*W J*_ and TC_*AT*_ (Figure 3A). The transitions between the open and closed-arm Y-shaped intermediates indicate that at [Mg^2+^] = 3 mM, AD populates conformations, which can either facilitate or hinder the formation of the ligand binding pocket.

**Figure 3:**
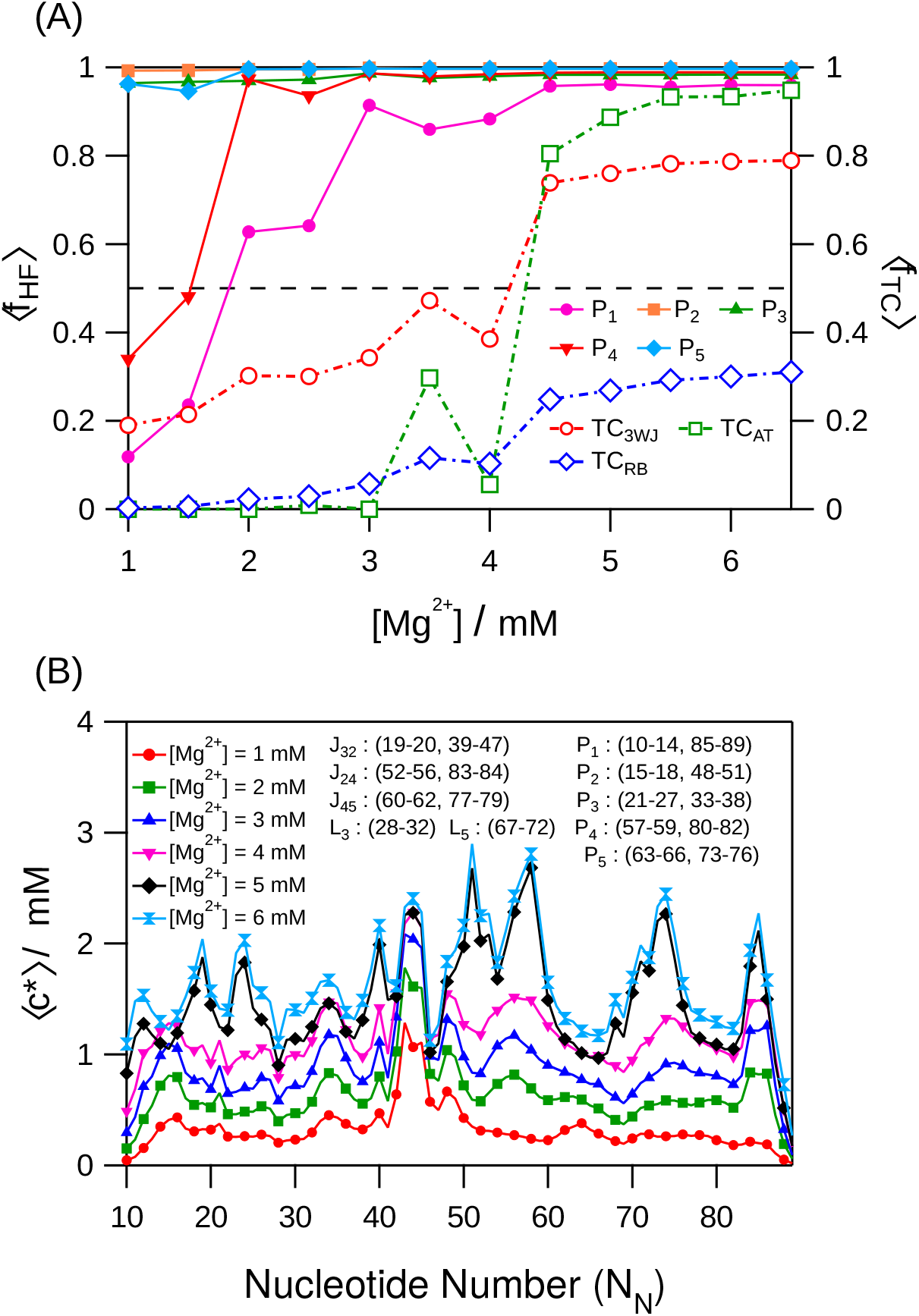
(A) Average fraction of helix formation (⟨ *f*_*HF*_ ⟩) and average fraction of tertiary contact formation (⟨ *f*_*TC*_ ⟩) plotted as a function of [Mg^2+^]. ⟨ *f*_*HF*_ ⟩ for helices P_1_ to P_5_ is plotted with solid magenta circle, orange square, green triangle, red inverted triangle, and cyan rhombus markers, respectively. Helices P_2_, P_3_ and P_5_ are stable for all [Mg^2+^]. Helices P_1_ and P_4_ are stable for [Mg^2+^] 2 mM. ⟨ *f*_*TC*_ ⟩ for TC_3*W J*_, TC_*AT*_ and TC_*RB*_ are plotted with hollow red circles, hollow green squares, and hollow blue rhombus markers, respectively. (B) The average local concentration of Mg^2+^ around individual P sites (*c*^*\**^) is plotted as a function of the nucleotide number (*N*_*N*_). The data for [Mg^2+^] = 1 to 6 mM are shown in solid red circle, green square, blue triangle, magenta inverted triangle, black rhombus, and cyan hourglass markers, respectively. The *N*_*N*_ range (starting from 10 as in the PDB^40^) for the helices (P’s), junctions (J’s), and loops (L’s) are in the annotation. For [Mg^2+^] = 1 mM, Mg^2+^ condenses around the J_32_ junction on the P_2*−*3_ arms. As [Mg^2+^] increases, it accumulates around the core and 3WJ regions leading to the AD folded structure.

### Stability of the Helical Arms

Out of all the five helices present in the AD, the P_1_ helix located at the basal region, which holds the two helical arms together, is thermodynamically the least stable helix and remains unfolded for [Mg^2+^] *<* 2 mM (Figure 1A, 2 and 3A). The lower stability of the P_1_ helix was also observed in previous studies.^25,95^ A single-molecule FRET experiment^18^ has shown that the P_1_ helix is highly dynamic and can exist in the unfolded form in the absence of Mg^2+^. Unfolded P_1_ helix leads to unzipping of the P_4_ helix in low [Mg^2+^] (*<* 2 mM) (Figure 2). The two arms of the AD retain their shape as the helices P_2_, P_3_, and P_5_ remain stable even in low [Mg^2+^] (= 1 mM) (Figure 3A). Monovalent cations ([K^+^] = 30 mM), which neutralize the RNA backbone charge, are sufficient to stabilize the P_2_, P_3_ and P_5_ secondary structures. However, the P_1_ helix is unfolded in the presence of K^+^ ions and low [Mg^2+^]. When [Mg^2+^] *≥* 2 mM, the P_1_ helix is stabilized, and it further stabilizes the P_2*−*3_ and P_4*−*5_ helical arms forming a Y-shaped structure (Figure 2). Fluorescence spectroscopy experiment^96^ has shown that at physiological [Mg^2+^], P_1_ helix orients the P_2*−*3_ and P_4*−*5_ arms forming a three-way-junction (3WJ).

### Site-Specific Mg^2+^ Binding Facilitates TC Formation

To understand the role of Mg^2+^ in TPP AD folding, we computed the average local Mg^2+^ concentration (⟨*c*\*⟩) around the AD at nucleotide resolution (Eq. 4). We found that Mg^2+^ are not randomly diffused along the RNA backbone but are bound to specific nucleotides and assist the formation of tertiary contacts.^4,56,69,97,98^ When [Mg^2+^] = 1 - 2 mM, Mg^2+^ are dominantly distributed around the nucleotides, *N*_*N*_ (nucleotide number) = A43 to A45, which belong to the J_32_ junction (Figure 3B). Due to the lack of Mg^2+^ in the core region and the three-way-junction (3WJ) (Figure 1A, 3B), the electrostatic repulsion between the P_2_, P_4_ helices and J_24_ junction (basal inter-arm motif) hinders the close approach of the P_2*−*3_ and P_4*−*5_ arms to form TCs and the folded state (Figure 2).

As [Mg^2+^] is increased to 3 - 4 mM, Mg^2+^ accumulates preferentially around the core region (cavity formed by the P_3_, J_32_, P_2_, J_24_, P_4_ and P_5_ motifs) and the 3WJ region (*N*_*N*_ = G33 - A35; A43 - A45; C48 - C49; U54 - A56; C57 - C58; C74 - G76; U14 - G16, A84 - G86, respectively) (Figure 3B, S2B). The condensed Mg^2+^ decrease the effective negative charge on the P sites of the basal inter-arm motif weakening the electrostatic repulsion between the P_2*−*3_ and P_4*−*5_ arms. As a result, the helical arms approach each other, and the Y-shaped intermediate state is populated (Figure 2).

When [Mg^2+^] is large (*≥* 5 mM), Mg^2+^ are condensed in the core and 3WJ regions with comparatively increased binding preference to the P_3_ (*N*_*N*_ = C23 - C24), J_32_ (*N*_*N*_ = G19 - U20 and U39 - A41 and A43 - A45), P_2_ (*N*_*N*_ = G17 - G18 and C48 - G51), J_24_ (*N*_*N*_ = C55 *≥*- A56), P_4_ (*N*_*N*_ = C57 - U59), P_5_ (*N*_*N*_ = C73 - A75), and L_5_ (*N*_*N*_ = U71 - G72) motifs (Figure 3B). The increased accumulation of Mg^2+^ on the helical arms and the J_24_ junction further weakens the inter-arm electrostatic repulsion and allows the arms to form TCs, which stabilizes the *holo*-form like folded structure even in the absence of the TPP ligand.

The nucleotides with high preferential binding of Mg^2+^, A43 and C74 to G76 are involved in the TPP ligand binding to the AD. A43 forms stacking interaction with the HMP group of the TPP ligand, and C74 - G76 bind to the pyrophosphate tail of TPP ligand bridged by Mg^2+^ (Figure S3).^40^ Mg^2+^ prefers binding to A43 even in low [Mg^2+^], whereas the ions show binding preference towards C74 - G76 when [Mg^2+^] *≥* 3 mM. The fact that Mg^2+^ prefers to bind to the nucleotides involved in TPP ligand recognition and binding illustrates that Mg^2+^ aids in the formation and stabilization of the scaffolding for ligand binding pocket.^18,40^ The SAXS experiments^42,43^ on *E. coli thiM* mRNA show that in high [Mg^2+^] (≈ 10 mM), TPP AD samples compact conformations similar to the *holo*-form like structures, whose *R*_*g*_ is greater than the ligand-bound structures by ≈ 2 Å. The experiments provide evidence that the T-loop (U39 - A47) region where the TPP ligand binds (Figure S2B) could be partially unfolded in the absence of the ligand.

### TC Formation is Cooperative

We computed the average fraction of TCs in TPP AD, ⟨ *f*_*TC*_ ⟩, to probe the cooperativity and hierarchy in TC formation with the variation in [Mg^2+^]. The two TCs located at the 3WJ (TC_3*W J*_) and the arm-tip of AD (TC_*AT*_) exhibit [Mg^2+^] dependent formation (Figure 3A). Both the TCs form cooperatively and show a sharp transition in the range 4 mM < [Mg^2+^] < 4.5 mM, which coincides with the second transition observed in both ⟨ *R*_*g*_ ⟩ and ⟨ *f*_*NC*_ ⟩ (Figure 1B) establishing that the compaction in AD dimensions is due to the formation of these TCs. The enhanced stability of the Y-shaped intermediates and the *holo*-form like state require the complete formation of these two TCs. The TC_3*W J*_ forms first and stabilizes the two helical arms, which subsequently approach each other to form TC_*AT*_ (Figure 3A). The tertiary structures do not form in the early stages of folding but during the AD assembly to the native-like state in the late stages of folding shows that TPP AD follows the quasi-hierarchical folding model. ^99^ TC_*RB*_ remains disrupted even in high [Mg^2+^]. We hypothesize that TPP ligand binding to the AD is essential to stabilize TC_*RB*_. The crystal structure of the TPP AD shows that the pyrophosphate domain of the TPP ligand interacts with G78 (J_45_) mediated by Mg^2+^ and water molecules (Figure S3C).

### Mg^2+^ Binding to Specific P Sites Decreases the Free Energy for TC Formation

To decipher the contribution of site-specific Mg^2+^ binding to the formation of a TC, we computed the difference in the stability of that TC due to Mg^2+^ binding and unbinding to the P site of *i*^*th*^ nucleotide, ΔΔ*G*_*TC*_(*i*) (Eq. 8).^70^ Negative value of ΔΔ*G*_*TC*_(*i*) for a TC indicates that Mg^2+^ binding to the *i*^*th*^ P site favors the formation of that TC. We computed ΔΔ*G*_*TC*_(*i*) for [Mg^2+^] = 4 mM, 4.5 mM and 5 mM, which correspond to the second transition in AD folding where TC formation is observed (Figure 1B and 3A). For [Mg^2+^] = 4 mM, conformations with both the TCs in the ruptured state are predominantly populated. With the increase in [Mg^2+^] (*≥* 4.5 mM), the AD dominantly populates conformations with the formed TCs (Figure S4).

Although, AD with the disrupted TC_3*WJ*_ is the most stable state at [Mg^2+^] = 4 mM, Mg^2+^ are already bound to the P sites located in the 3WJ region (*N*_*N*_ = G19, C50 - A53, A56 - C58, A85) (Figure 3A,B). The bound Mg^2+^ significantly favor the TC_3*W J*_ formation as ΔΔ*G*_*TC*_(*i*) < -0.5 *k*_B_*T* for *i* belonging to the 3WJ nucleotides (Figure 4A). With the increase in [Mg^2+^] (*≥* 4.5 mM), the probability of TC_3*W J*_ formation increased (Figure 3A). However, the ΔΔ*G*_*TC*_(*i*) values remain almost unchanged (Figure 4A), but the ⟨*c*\*⟩ around the *N*_*N*_ forming the TC_3*W J*_ shows a significant increase with the increase in [Mg^2+^] (*≥* 4.5 mM) (Figure 3B, S5). We hypothesize that at [Mg^2+^] = 4 mM, as lower number of Mg^2+^ accumulate in the 3WJ region, the backbone electrostatic repulsion may still prevail in the region and hinder the TC_3*W J*_ formation. With increasing [Mg^2+^] (*≥* 4.5 mM), sufficient number of Mg^2+^ accumulate in the 3WJ region (Figure 3B) and diminish the backbone electrostatic repulsion to stabilize the TC_3*W J*_ (Figure 3A, 4A).

**Figure 4:**
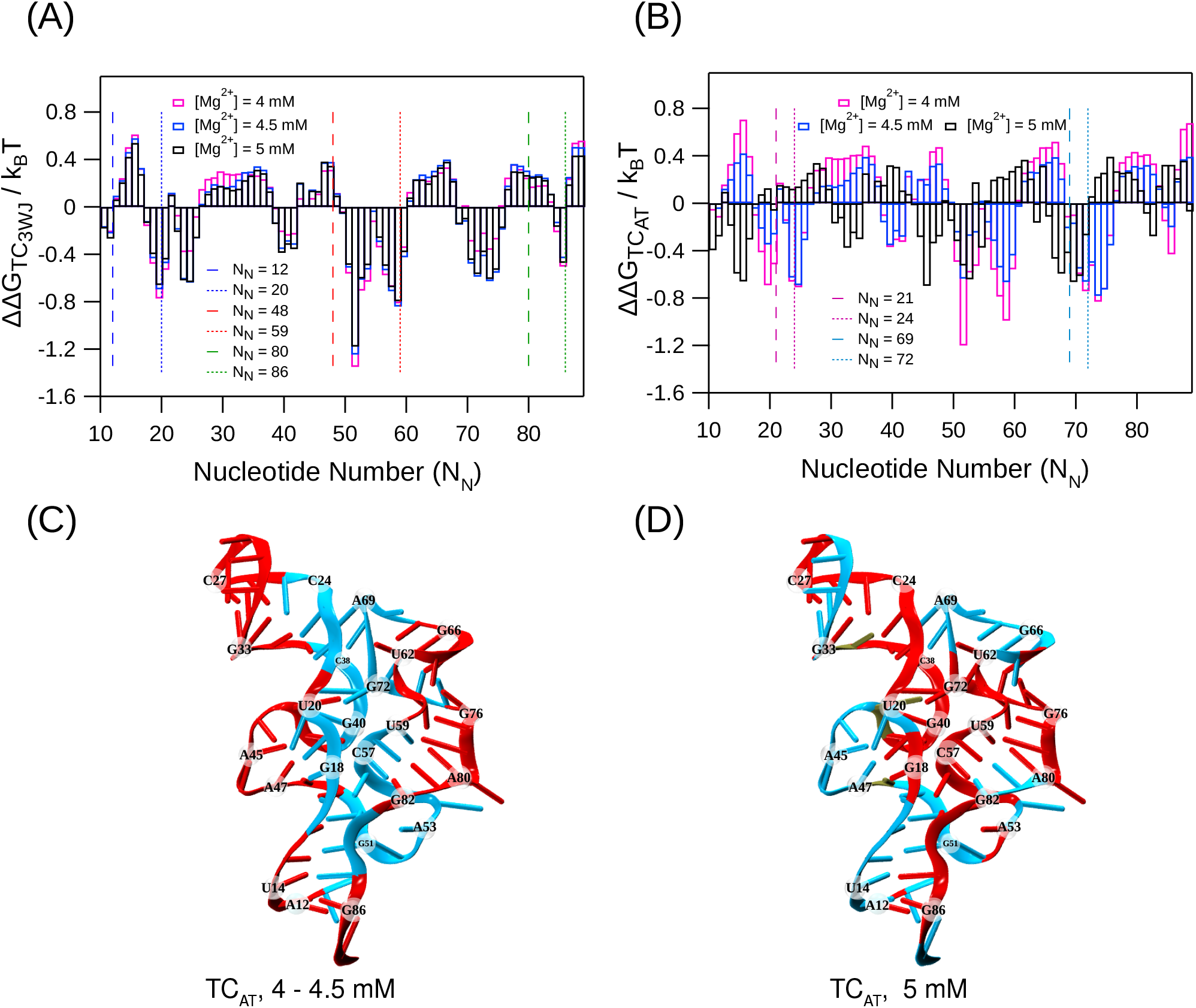
ΔΔ*G*_*TC*_ for (A) TC_3*W J*_ and (B) TC_*AT*_ are plotted as a function of *N*_*N*_ for [Mg^2+^] = 4 mM, 4.5 mM and 5 mM in magenta, blue, and black bars, respectively. The dashed and dotted lines mark the range of nucleotides involved in the formation of a particular TC. Nucleotides, which facilitate (ΔΔ*G*_*TC*_(*i*) < 0) and hinder (ΔΔ*G*_*TC*_(*i*) > 0) the formation of TC_*AT*_ formation upon Mg^2+^ binding are shown in cyan and red colored cartoon representation for (C) [Mg^2+^] = 4 and 4.5 mM (D) [Mg^2+^] = 5 mM.

During the second transition in AD folding ([Mg^2+^] = 4 - 4.5 mM), TC_*AT*_ formation is favored by Mg^2+^ binding to the nucleotides located at the interior region mainly to the nucleotides located at both 3WJ (*N*_*N*_ = G19 - U20, C50 - A53, A56 - U59 and A85) and arm-tip (*N*_*N*_ = C22 - C23, A70 - G72) regions of the AD (Figure 4B,C). The lower value of 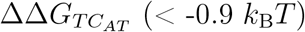 due to Mg^2+^ binding to the 3WJ nucleotides (*N*_*N*_ = G51 - C58) compared to the arm-tip nucleotides, indicates that Mg^2+^ binding in the 3WJ region can expedite the TC_*AT*_ formation, located at a distant site through allosteric interactions. At [Mg^2+^] = 4.5 mM, AD dominantly populates the conformations with formed TC_*AT*_ (Figure 3A). Mg^2+^ binding to both 3WJ (*N*_*N*_ = G51, C58) and arm-tip (*N*_*N*_ = C23 - C24, A70 - G72) nucleotides almost equally contribute towards TC_*AT*_ formation. After the second transition ([Mg^2+^] = 5 mM), TC_*AT*_ formation is governed by the Mg^2+^ binding to the nucleotides located at the periphery of AD (Figure 4B,D). Most of these peripheral nucleotides, *N*_*N*_ = C30 - U32 (L_3_), A44 - A47 (J_32_) and A67 - U71 (L_5_), belong to motifs devoid of any three-dimensional structures. The favorable free energy for TC_*AT*_ formation due to Mg^2+^ binding to the peripheral nucleotides can probably lead to the proper orientation of the junction and loop region so that AD can fold to its *holo*-form like conformation.

ΔΔ*G*_*TC*_(*i*) data reveals that the nucleotides located near the 3WJ region facilitate the AD folding to its *holo*-form like state by stabilizing both the TCs. The stabilization of TCs due to the site-specific Mg^2+^ binding to the nucleotides in the 3WJ region irrespective of [Mg^2+^] establishes that Mg^2+^ binds to specific nucleotides of an RNA molecule that drive and stabilize the TC formation leading to the RNA folded state.^70^

### TPP AD Folds Through Slow and Fast Folding Pathways

We spawned 60 folding trajectories to study the AD folding kinetics in the absence of the TPP ligand starting from different unfolded conformations at *T* = 310 K and [Mg^2+^] = 6.5 mM. The initial unfolded conformations are taken from the [Mg^2+^] = 1.0 mM simulation, where the unfolded state is the most stable state (Figure 2). Upon initiating folding, we find that the AD folds through fast and slow folding pathways. We labeled trajectories where TPP AD folds in less than 0.5 *µ*s as fast folding pathways and the other trajectories as slow folding pathways. 16 out of the 60 spawned folding trajectories follow the fast folding pathway and the rest fold through the slow folding pathway. In the slow folding pathway, an intermediate, which acts as a kinetic trap, is populated, leading to a longer folding time (Figure 5B,D).

**Figure 5:**
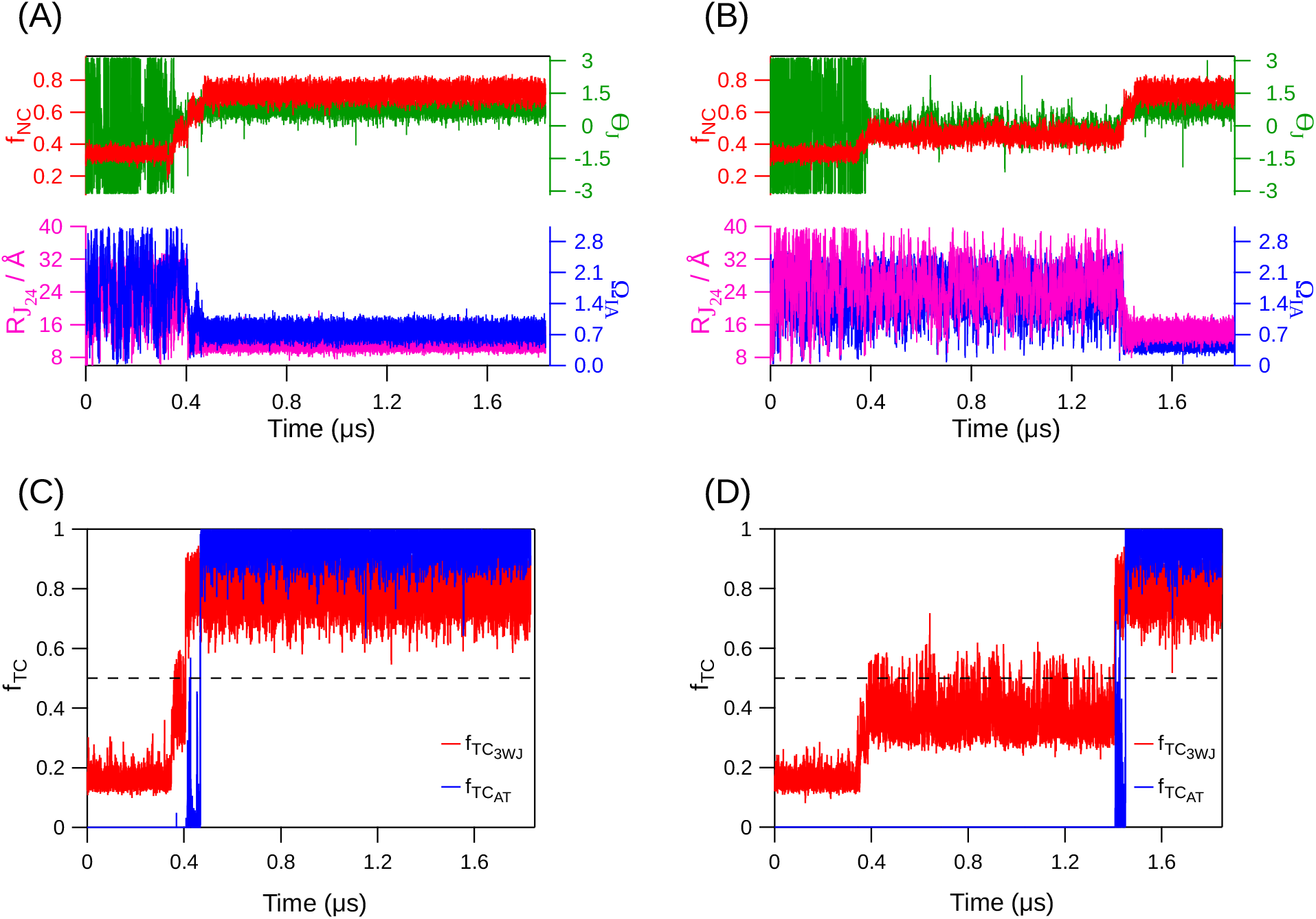
Fast and slow folding TPP AD pathways at *T* = 310 K and [Mg^2+^] = 6.5 mM. Fraction of native contacts (*f*_*NC*_), dihedral angle between the two strands of J_24_ motif (Θ_*J*_), inter-helical arm angle (Ω_*IA*_), and length of the J_24_ motif 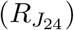 are plotted for (A) fast and (B) slow folding pathways with red, green, blue, and magenta solid lines, respectively. For both the pathways, Ω_*IA*_ and 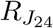 exhibit large fluctuations till AD folds to the native state. Fluctuations in Θ_*J*_ decrease after the formation of the *P*_1_ helix and fluctuate around 0.75 rad as the AD folds to the native state. Fraction of TC formation (*f*_*TC*_) as a function of time is plotted for (C) fast and (D) slow folding pathways. 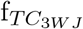 and 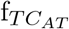 for both fast and slow folding trajectories are shown in red and blue solid lines, respectively. The TCs form during the late stages of folding and follow a hierarchical trend. In the slow folding trajectory, the kinetic trap is observed in the time interval 0.39 *µ*s to 1.41 *µ*s, where Θ_*J*_ fluctuates around -0.1 rad and *f*_*TC*_ fluctuates between 0.4 - 0.5 rad.

The helices P_2_, P_3_ and P_5_ are stable in the unfolded state at [Mg^2+^] = 6.5 mM (Figure 3A). The Mg^2+^ mediated formation of the least stable switch helix P_1_ is essential for the formation of 3WJ and open-arm Y-shaped intermediate for the folding of the AD in the absence of ligand. The initial barrier that AD has to overcome to fold is the relative orientation of the two helical arms P_2*−*3_ and P_4*−*5_ with respect to each other. In the early stages of folding, the arms fluctuate relative to each other as inferred from the inter-arm angle, Ω_*IA*_ (see Methods) (Figure 5A,B and 6A,A^*1*^). Stochastic fluctuations rotate the P_4*−*5_ arm around the J_24_ junction as an axis to align it parallel to the P_2*−*3_ arm (Figure 6B,B^*1*^). The parallel alignment of both the arms also facilitates the formation of P_1_ helix at the base (Figure 6C_1_,C^*1*^). The fluctuations in Ω_*IA*_ show that the parallel orientation of the helical arms is necessary but not sufficient for the AD to proceed to fold to its native state (Figure 5A,B).

**Figure 6:**
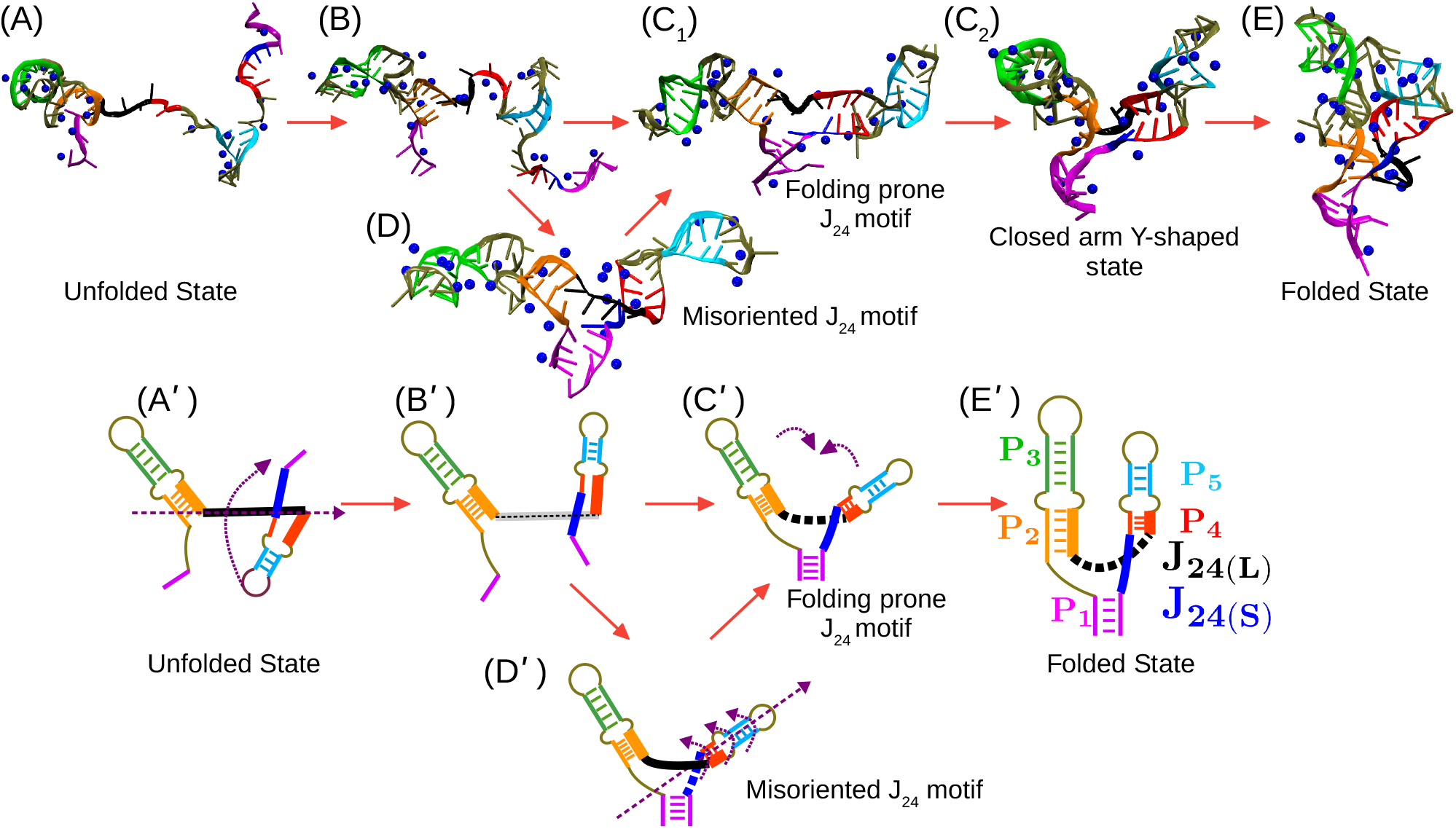
Intermediates populated during AD folding are shown in cartoon and schematic representations. Helices P_1_ to P_5_ are shown in magenta, orange, green, red, and cyan, respectively. The blue spheres are Mg^2+^ within 5 Å from the RNA. J_24(*L*)_ and J_24(*S*)_ are shown in black and blue lines, respectively. The rotational axis, the direction of rotation, and helical movement are shown as purple dashed and dotted arrows, respectively. The shaded black dot line in (B^*1*^) represents that the relative orientation between J_24(*L*)_ and J_24(*S*)_ motifs cannot be clearly demarcated as the P_4_ helix is not formed at that stage of folding. The red arrows show the progression of folding events with time. (A) and (A^*1*^) Helical-arms are not in parallel orientation with respect to each other. The P_4*−*5_ arm rotates around the J_24(*L*)_ motif to attain the folding prone conformation (B) and (B′). The AD at stage (B) and (B′) has neither adopted the folding prone conformations (C_1,2_) and (C′) of J_24(*L*)_ motif nor the misoriented (D) and (D′). In (C_1_) and (C′), the P_2_-J_24(*L*)_-P_4_ and P_1_-P_2_/P_4_-J_24(*S*)_-P_1_ motifs (see SI for structural details) do not cross each other and the AD folds to form closed-arm Y-shaped conformation (C_2_), which eventually leads to the *holo*-form like folded state (E) and (E′). (D) and (D′) The AD is stuck in the kinetic trap and to escape from the trap, J_24(*L*)_ motif, P_4_-P_5_ helices rotate around the P_4*−*5_ arm as an axis to attain the folding prone structure.

### Misaligned Orientation of P_2_-J_24(L)_-P_4_ and P_1_-P_2_/P_4_-J_24(S)_-P_1_ Motifs Leads to a Kinetic Trap

The kinetic trap in the slow folding pathway is due to the population of a conformation where although both the arms are aligned parallel to each other, their orientation is misaligned, and this conformation with a lifetime greater than 1 *µ*s acts as a kinetic trap (Figure 5B,D and 6D,D′). In the kinetic trap, the nucleotides G51 - C57 (P_2_-J_24(*L*)_-P_4_ motif) are misaligned with respect to the nucleotides 14U - 15C and G82 - A85 (P_1_-P_2_/P_4_-J_24(*S*)_-P_1_ motif). We quantified this misalignment using the dihedral angle Θ_*J*_ (see Methods). For both the slow and fast folding pathways, Θ_*J*_ initially fluctuates due to the unfolded P_1_ and P_4_ helices. After the formation of the helices, the magnitude of the fluctuations in Θ_*J*_ decreased. If the AD is stuck in the kinetic trap, Θ_*J*_ fluctuates around -0.1 rad (Figure 5B). The misaligned arms prevent the formation of pre-organized 3WJ, which facilitates the formation of the TC_*AT*_ tertiary structure. To escape from the kinetic trap, J_24(*L*)_, P_4_ and P_5_ rotate around the P_4*−*5_ arm (Figure 6C_1_,D,C′,D′). After escaping from the kinetic trap, a U-shaped loop is formed by the longer part of the J_24_ junction (J_24(*L*)_) joining the P_2_ and P_4_ helices, which allows the helical arms to come closer to acquire native-like folded state competent to form the ligand-binding pocket (Figure 6C_2_,E,E′). The loop-formation by the folded J_24(*L*)_ junction pulls the helical arms together leading to the formation of TC_*AT*_ and *holo*-form like folded state. The inter-arm angle, Ω_*IA*_ and 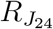 show large fluctuations when the AD is unfolded, and fluctuations subside only after the AD is completely folded.

### Mg^2+^ Binding to Specific P Sites Facilitates Folding to the Native State

To probe the effect of Mg^2+^ on TPP AD folding, we computed *c*\* around the P sites of nucleotides involved in the formation of TCs (P_*TC*_ sites) (Figure S6 and S7) and *f*_*TC*_ as a function of time (*t*_*kin*_) (Figure 5C,D). In the slow folding pathway, during the early stages of folding (*t*_*kin*_ *⪅* 0.39 *µ*s), *c*\* varies between 0 - 1 mM for all the P_*TC*_ sites and none of the TCs are formed. After the population of the kinetic trap due to the misalignment of the two arms (0.39 *µ*s < *t*_*kin*_ < 1.41 *µ*s), TC_3*W J*_ fluctuates between states with *f*_*TC*_ = 0.4 to 0.5, as the AD attempts to escape from the kinetic trap (Figure 5D). The *f*_*TC*_ plot shows hierarchy in the formation of the TCs and the formation of TC_3*W J*_ is a prerequisite for the formation of TC_*AT*_ (Figure 5C,D). Both the TCs form after native-like reorganization of the J_24_ motifs at *t*_*kin*_ > 1.41 *µ*s. Similar time-dependent trends are also observed in *c*\* for fast-folding pathways, where the formation of the TCs is facilitated by Mg^2+^ accumulation around the P_*TC*_ sites. Hierarchy in TC formation where TC_3*W J*_ anchors the formation of TC_*AT*_ is also observed in the fast folding pathways as well. Both thermodynamics and kinetics folding studies confirm that AD folding follows a quasi-hierarchical model, where TCs are not formed during the early stages of folding, and they form only during the AD assembly in the late stages of folding.

## Conclusion

In this study, we showed that TPP AD populates intermediates when its folding is induced by Mg^2+^ in agreement with the experiments.^12^ We found that at physiological [Mg^2+^] (≈ 4 mM), AD populated a closed-arm Y-shaped and a *holo*-form like folded conformations. The *holo*-form like conformation is not the global minimum in the FES at around physiological [Mg^2+^] but becomes the global minimum in higher [Mg^2+^] (*≥* 5 mM) even in the absence of the TPP ligand. These results indicate that *holo*-form like riboswitch conformations are only marginally stable at physiological conditions in the absence of their cognate ligands. Therefore at physiological conditions, TPP-ligand binding to the AD is indispensable for gene regulation and further supports the hypothesis that gene regulation by riboswitches is under kinetic control. The population of *holo*-form like conformations in the absence of cognate ligand but in high [Mg^2+^] is also observed in preQ_1_, Fluoride and SAM-I riboswitch.^28,44,45,100,101^ Experiments on preQ_1_ riboswitch have shown that Mg^2+^ can further shift the aptamer-ligand sensing mechanism from induced-fit to conformation selection.^45^ The unique ability of Mg^2+^ to stabilize the *holo*-form like AD conformations in the absence of the cognate ligand poses an interesting unanswered question whether only Mg^2+^ can lead to the population of *holo*-form like structures in the AD of all riboswitches without their cognate ligands and what is the optimum Mg^2+^ concentration required?

## Acknowledgement

A part of this work is funded by the grant to G.R. by the National Supercomputing Mission (MeitY/R&D/HPC/2(1)/2014). S.K. acknowledges research fellowship from the Indian Institute of Science, Bangalore. We acknowledge National Supercomputing Mission (NSM) for providing computing resources of “PARAM Brahma” at IISER Pune, which is implemented by C-DAC and supported by the Ministry of Electronics and Information Technology (MeitY) and Department of Science and Technology (DST), Government of India.

## Supporting Information Available

Structural details of P_2_-J_24(L)_-P_4_ and P_1_-P_2_/P_4_-J_24(S)_-P_1_ motifs is provided in supporting information. Figure S1: Native contact map of AD; Figure S2: 3D structure of the AD with highlighted TC_3*W J*_ and TC_*AT*_ forming nucleotides and 2D schematic structure of the AD with definition of parameters used to characterize the kinetic results; Figure S3: Atomistic representation of RNA residues involved in the TPP ligand binding; Figure S4: Time profile of *f*_*TC*_ for both TC_3*W J*_ and TC_*AT*_ showing the formation of TCs at [Mg^2+^] = 4 to 5 mM; Figure S5: ⟨ *c*\* ⟩ for [Mg^2+^] = 4 to 5 mM; Figure S6-S7: Time profile of *c*\* at TC forming nucleotide resolution for fast and slow folding trajectories; Table S1-S6: Various parameters used to model the TPP AD using TIS model is given in the tabular format in the supporting information.

## For Table of Contents Use Only

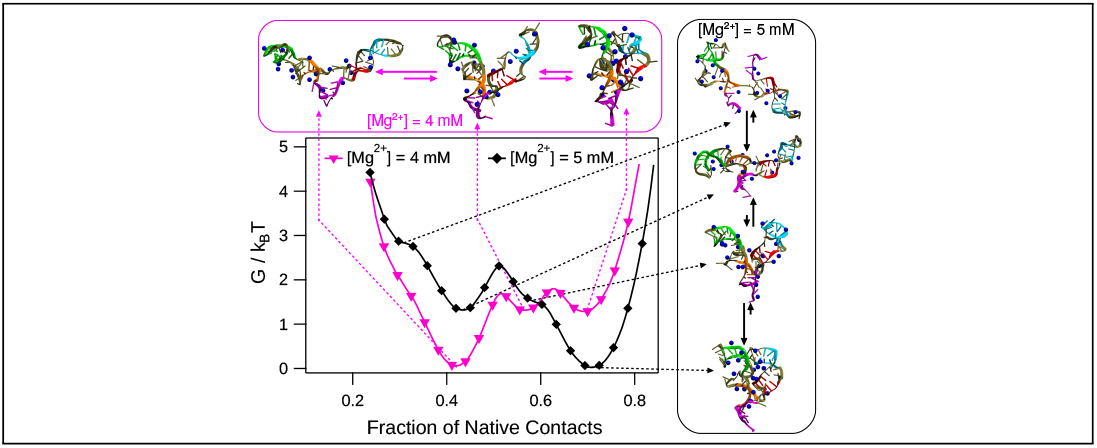

