## Supporting Information for "Riboswitch Folds to *Holo*-Form Like Structure Even in the Absence of Cognate Ligand at High Mg^2+^ Concentration"

**Structural details of $P_2$-$J_{24(L)}$-$P_4$ and $P_1$-$P_2$/$P_4$-$J_{24(S)}$-$P_1$ Motifs:** The $P_2$-$J_{24(L)}$-$P_4$ motif is composed of the longer part of the $J_{24}$ junction (U52 to A56), $J_{24(L)}$(shown in black color in Figure 6), connecting the $P_2$ and $P_4$ helices through G51 and C57 whereas the $P_4$-$J_{24(S)}$-$P_1$ motif is composed of the shorter part of the $J_{24}$ junction ($N_N$ = G83 - A84), $J_{24(S)}$ (shown in blue color in Figure 6) connecting $P_4$ and $P_1$ helices through G82 and A85. To ensure that the aptamer folds to its native-state, the $P_2$-$J_{24(L)}$-$P_4$ motif should never cross-over the $P_1$-$P_2$/$P_4$-$J_{24(S)}$-$P_1$ motif.

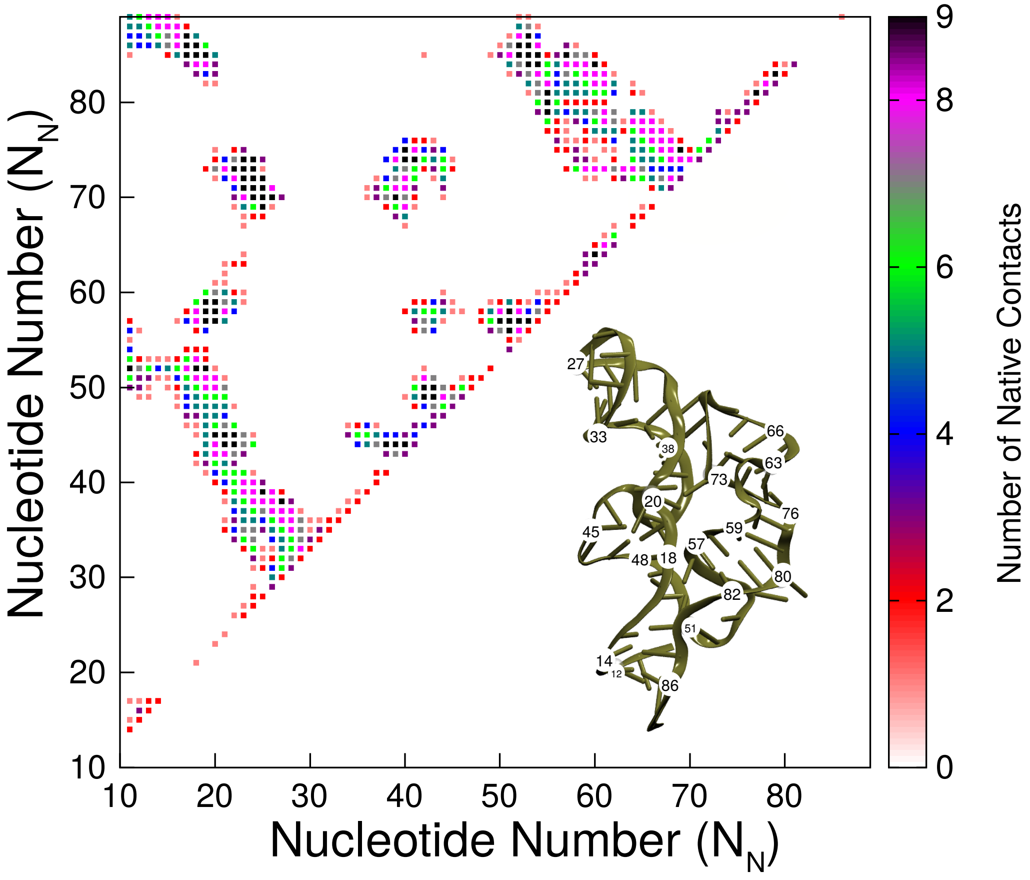


Figure S1: Native contact map of the TPP AD at the nucleotide resolution, and the native structure is shown in the inset.

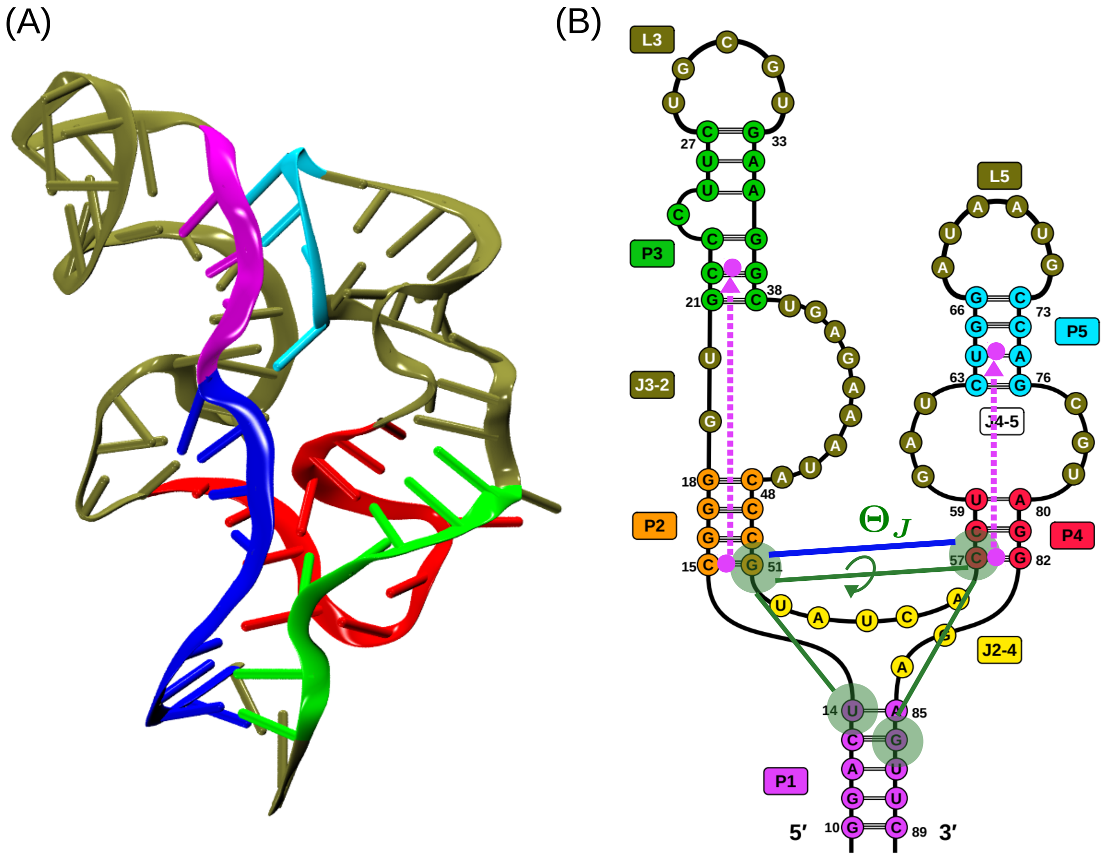


Figure S2: (A) 3D structure of the TPP AD is shown in cartoon representation (PDB: 2GDI).[1] $TC_{3WJ}$ forming nucleotides, $N_N$ = A12 - U20, C48 - U59 and A80 - G86 are shown in blue, red and green, respectively. $TC_{AT}$ forming nucleotides, $N_N$ = G21 - C24 and A69 - G72 are shown in magenta and cyan, respectively. (B) 2D Schematic of the TPP AD is shown with the nucleotides numbered. Helices $P_1$ to $P_5$ are shown in magenta, orange, green, red, and cyan, respectively. Helical junctions ($J_{32}$ and $J_{45}$) and loops ($L_3$ and $L_5$) are shown in tan. Junction $J_{24}$ is shown in yellow. $\Omega_{IA}$ is the angle between vectors shown in dotted magenta lines. $\Theta_J$ is the dihedral angle between S sites of the nucleotides U14, G51, C57 and G86 shown in shaded green circle connected with green solid lines. The end-to-end distance of $J_{24(L)}$ motif ($R_{J_{24}}$) is the distance between P sites of the nucleotides G51 - C57, and is shown by the solid blue line.

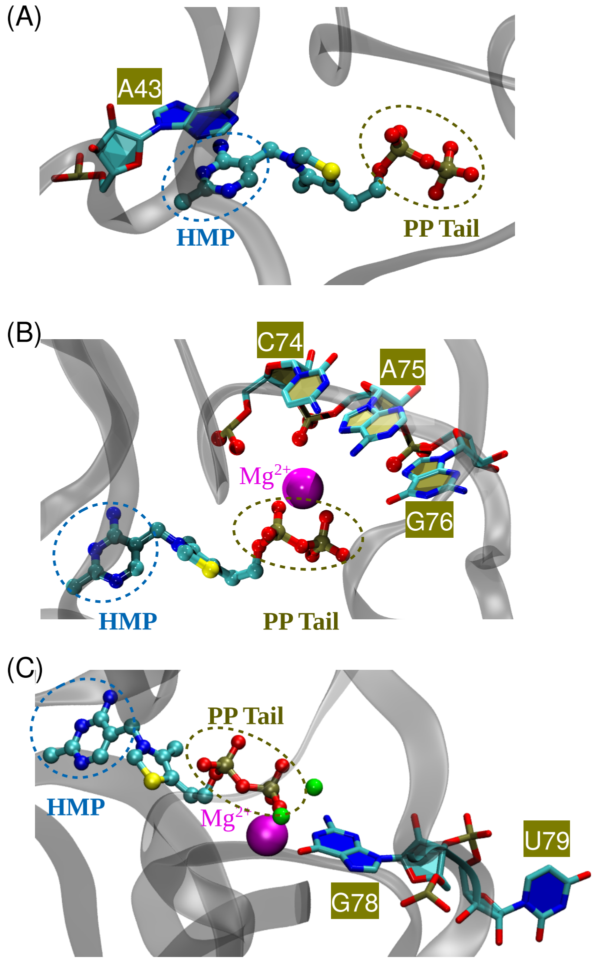


Figure S3: TPP ligand interaction with the TPP AD (PDB ID: 2GDI). TPP AD backbone is shown in transparent grey cartoon and TPP ligand in ball-stick representation. Atoms C, N, O, P, S and Mg are shown in cyan, blue, red, tan, yellow, and magenta colored beads, respectively. The O atom of water molecules are shown as green colored beads. (A) HMP ring of TPP ligand interacts with the base of A43 through $\pi$-$\pi$ stacking. (B) Pyrophosphate tail of the ligand interacts with the phosphate group O atoms of the nucleotides, $N_N$ = C74 - G76, mediated through $Mg^{2+}$. (C) Pyrophosphate tail of the ligand interacts with the atoms of the nucleotide G78 mediated through $Mg^{2+}$ and water molecules.

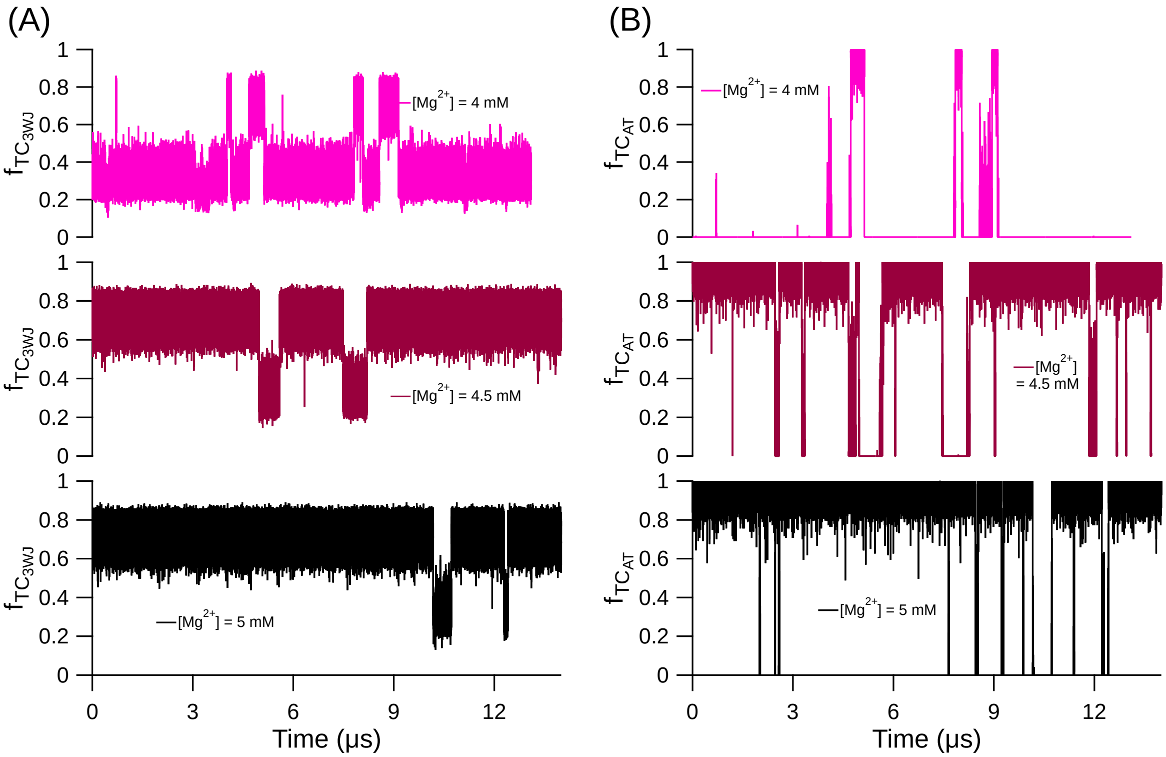


Figure S4: Fraction of tertiary contacts ($f_{TC}$) for TCs are plotted as a function of simulation time for [$Mg^{2+}$] = 4 mM, 4.5 mM and 5 mM for (A) $TC_{3WJ}$ and (B) $TC_{AT}$. The AD dominantly populates conformations with formed TCs for [$Mg^{2+}$] $\geq$ 4.5 mM.

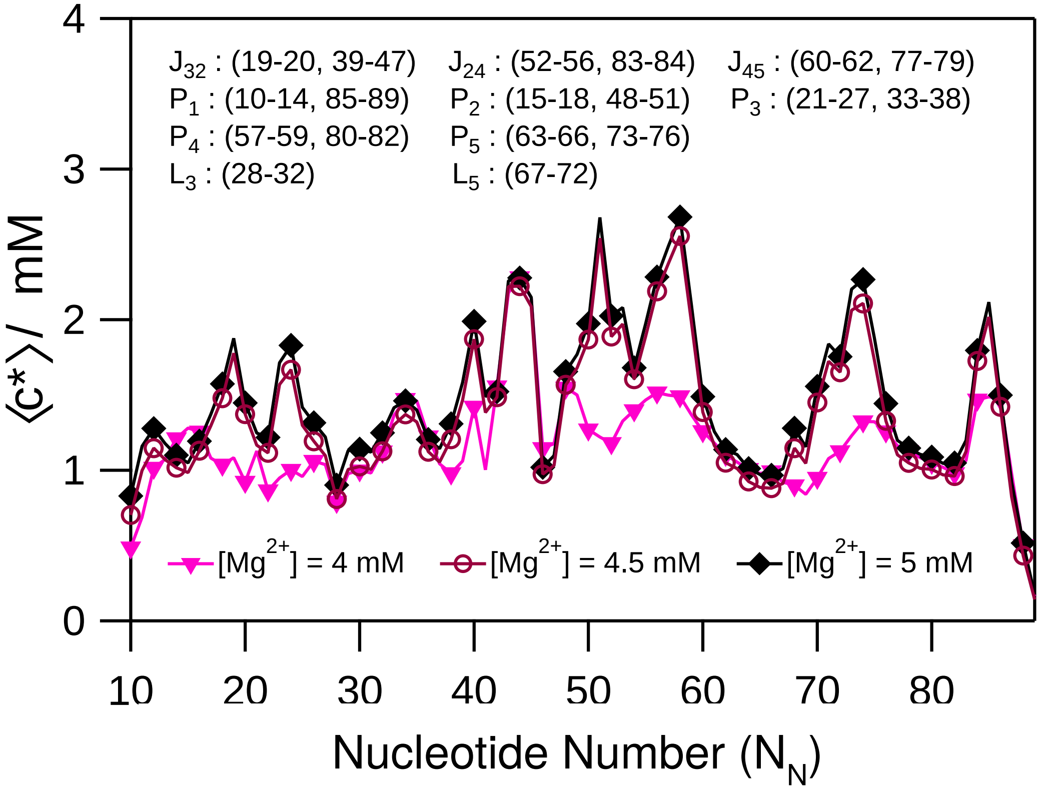


Figure S5: The average local concentration of $Mg^{2+}$ ($\langle c^{*}\rangle$) around the P sites is plotted as a function of the nucleotide number ($N_N$). Data for $[Mg^{2+}]$ = 4 to 5 mM are shown in solid magenta inverted triangle, hollow maroon circle, and solid black rhombus markers, respectively. The $N_N$ range (starting from 10 as in the PDB ID: 2GDI) for the helices (P's), junctions (J's), and loops (L's) are shown in the annotation. For $[Mg^{2+}] \geq 4.5$ mM, $Mg^{2+}$ shows enhanced preference to bind to the nucleotides located in the 3WJ region.

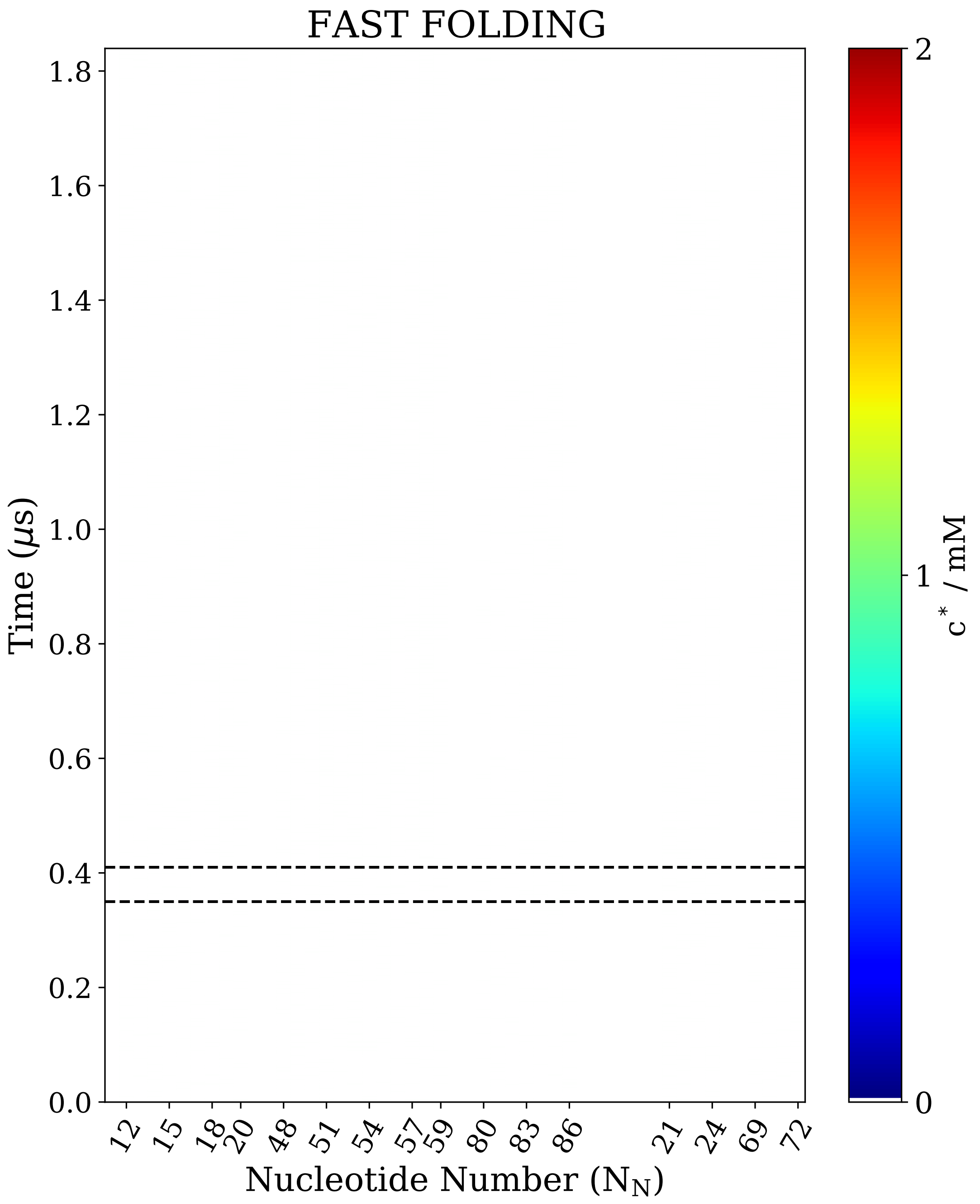


Figure S6: The local concentration of $Mg^{2+}$ around P sites ($c^{*}$) of the nucleotides involved in the $TC_{3WJ}$ and $TC_{AT}$ formation plotted as a function of the folding time for the fast folding trajectory. Nucleotides with $N_N$ = A12 - U20, C48 - U59 and A80 - G86 are involved in the $TC_{3WJ}$ formation. Nucleotides with $N_N$ = G21 - C24 and A69 - G72 are involved in the $TC_{AT}$ formation. The two dashed lines are drawn at the folding time $\approx 0.35$ and $\approx 0.41$ $\mu$s to demarcate the starting of size compaction and commencement of AD folding.

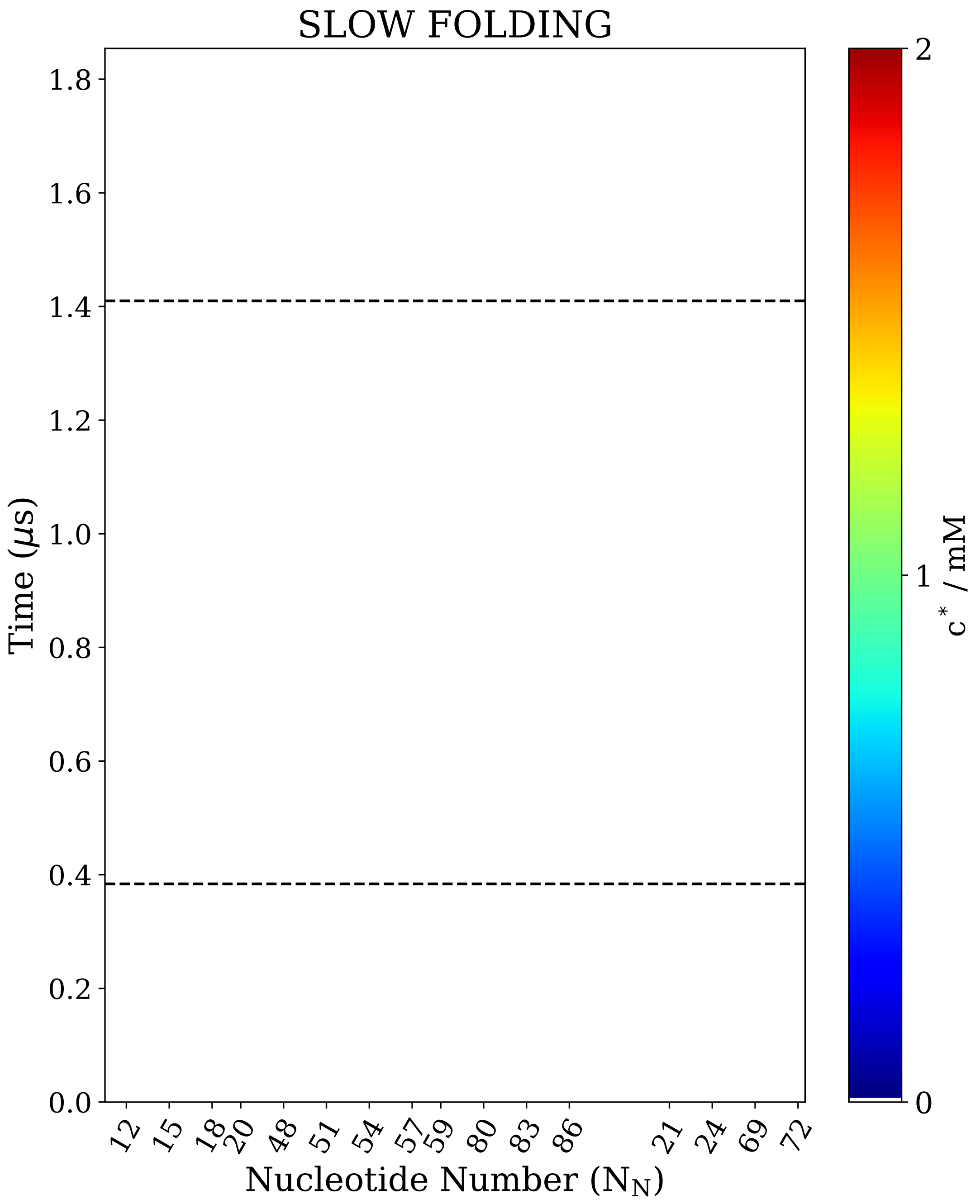


Figure S7: $c^{*}$ for the P sites of the nucleotides involved in the $TC_{3WJ}$ and $TC_{AT}$ formation plotted as a function of the folding time for the slow folding trajectory. The two dashed lines are drawn at the folding time $\approx 0.39$ and $\approx 1.41$ $\mu$s to demarcate the starting of size compaction and commencement of AD folding.

**Table S1: Parameters for the hydrogen bonding potential ($U_{HB}$).[2] The potential is given by** $U_{HB} = U_{HB}^0 \ \exp(-u)$**, where** $u = \Bigg[5(r - r_0)^2 + 1.5\Bigg\{(\theta_1 - \theta_{1,0})^2 + (\theta_2 - \theta_{2,0})^2)\Bigg\} + 0.15\Bigg\{(\psi - \psi_0)^2 + (\psi_1 - \psi_{1,0})^2 + (\psi_2 - \psi_{2,0})^2\Bigg\}\Bigg]$

**(a) Canonical hydrogen bond parameter.**

Parameters are derived from coarse-grained structure of ideal A-form RNA helix.[3]

| base pair | $r_0$ (Å) | $\theta_{1,0}$ | $\theta_{2,0}$ | $\psi_0$ | $\psi_{1,0}$ | $\psi_{2,0}$ |
|---|---|---|---|---|---|---|
| A−U | 5.8815 | 2.7283 | 2.5117 | 1.2559 | 0.9545 | 1.1747 |
| C−G | 5.655 | 2.4837 | 2.823 | 1.3902 | 1.2174 | 0.7619 |

**(b) Nucleotide number of base-pairs forming canonical hydrogen bond in TPP AD.**

| nucleotide($B_1$)–nucleotide($B_2$), (#H-bonds) | | | |
|---|---|---|---|
| 11G–88U,(3) | 12A–87U,(2) | 13C–86G,(3) | 14U–85A,(2) |
| 15C–51G,(3) | 16G–50C,(3) | 17G–49C,(3) | 18G–48C,(3) |
| 19G–48C,(1) | 20U–45A,(2) | 21G–71U,(1) | 21G–38C,(3) |
| 22C–37G,(3) | 23C–36G,(3) | 24C–37G,(1) | 25U–35A,(2) |
| 26U–34A,(2) | 27C–33G,(3) | 39U–43A,(2) | 57C–82G,(3) |
| 58C–81G,(3) | 59U–80A,(2) | 63C–76G,(3) | 64U–75A,(2) |
| 65G–74C,(3) | 66G–73C,(3) | | |

**Table S2: Nucleotide number of base-pairs forming sugar-sugar hydrogen bond in TPP AD.**

Parameters are derived from coarse-grained structure of TPP AD.

| $S_1$–$S_2$ (#H-bonds) | $r_0$ | $\theta_{1,0}$ | $\theta_{2,0}$ | $\psi_0$ | $\psi_{1,0}$ | $\psi_{2,0}$ |
|---|---|---|---|---|---|---|
| 43–20 | 5.8148 | 1.6863 | 2.1264 | -0.5485 | -1.3762 | -1.4883 |
| 47–45 | 6.3517 | 2.8883 | 1.6273 | 2.9975 | 0.1371 | 0.6872 |
| 48–41 | 6.5557 | 1.5202 | 1.8685 | 2.7877 | 1.2949 | 0.5829 |
| 56–49 | 6.4752 | 1.8749 | 1.5685 | 2.3463 | 2.0752 | 0.6281 |
| 57–18(2) | 5.3056 | 2.1098 | 1.6158 | 0.0052 | 1.0624 | 0.7806 |
| 70–22(2) | 6.4042 | 1.6829 | 1.2534 | 1.8795 | 0.3324 | 1.1083 |
| 71–21 | 8.1033 | 1.7777 | 1.2358 | 2.0726 | 0.865 | 0.5901 |
| 73–39 | 5.3528 | 1.7269 | 2.1868 | -0.4798 | 1.885 | 1.2782 |
| 84–16 | 6.6959 | 1.5392 | 1.5158 | 2.4942 | 1.4802 | 0.5686 |

**Table S3: Nucleotide number of base-pairs forming base-sugar hydrogen bond in TPP AD.**

Parameters are derived from coarse-grained structure of TPP AD.

| B-S (#H-bonds) | $r_0$ | $\theta_{1,0}$ | $\theta_{2,0}$ | $\psi_0$ | $\psi_{1,0}$ | $\psi_{2,0}$ |
|---|---|---|---|---|---|---|
| 19–42 | 8.2726 | 1.9843 | 1.7582 | -1.4425 | 1.6779 | 0.5782 |
| 21–43 | 6.1087 | 1.8338 | 1.832 | 1.3902 | -2.304 | -2.6195 |
| 24–37 | 6.764 | 2.6934 | 2.0401 | 2.6167 | 0.5445 | 0.6069 |
| 40–43(3) | 6.5334 | 2.5245 | 2.4729 | 3.0255 | 2.1503 | 2.108 |
| 41–56 | 6.9633 | 2.7493 | 1.1843 | 1.954 | -2.7763 | 2.0552 |
| 41–48 | 6.7042 | 1.1644 | 1.4948 | 1.5776 | 1.7342 | 0.5373 |
| 42–40(2) | 6.5422 | 1.6012 | 1.8075 | -1.2677 | -1.5126 | 0.44 |
| 49–56 | 7.9214 | 0.9557 | 2.0616 | 1.3673 | 1.5841 | 1.4851 |
| 56–83 | 7.575 | 2.6366 | 2.0482 | -2.5279 | -1.1623 | -0.1688 |
| 56–17 | 7.1082 | 2.4489 | 1.7078 | -1.5324 | 1.7826 | 0.4694 |
| 69–38 | 6.6497 | 2.5251 | 2.6394 | -0.3235 | -1.1984 | 1.7281 |
| 70–68 | 6.6041 | 1.4606 | 1.7664 | -1.5176 | -1.3012 | 0.639 |
| 70–22 | 6.8307 | 1.1092 | 1.8255 | 0.7391 | 1.5476 | 0.5963 |
| 70–38 | 6.4275 | 2.9925 | 2.4065 | 2.987 | 2.8653 | 0.8047 |
| 72–19 | 7.3275 | 2.1645 | 0.8823 | -3.0529 | 1.2833 | 1.5589 |
| 83–53 | 7.0864 | 2.4582 | 2.1718 | -2.37 | 0.047 | -1.0722 |
| 84–50 | 6.7848 | 2.5701 | 1.4575 | -1.5704 | 1.4032 | 0.3005 |

**Table S4: Nucleotide number of base-pairs forming phosphate-sugar hydrogen bond in TPP AD.**

Parameters are derived from coarse-grained structure of TPP AD.

| P–S | $r_0$ | $\theta_{1,0}$ | $\theta_{2,0}$ | $\psi_0$ | $\psi_{1,0}$ | $\psi_{2,0}$ |
|---|---|---|---|---|---|---|
| 24–69 | 6.0025 | 1.1795 | 1.8144 | 2.8922 | -2.8567 | 0.7822 |

**Table S5: Nucleotide number of base-pairs forming non-canonical hydrogen bond in TPP AD.**

Parameters are derived from coarse-grained structure of TPP AD (PDB: 2GDI).

| $B_1-B_2$(#H-bonds) | $r_0$ | $\theta_{1,0}$ | $\theta_{2,0}$ | $\psi_0$ | $\psi_{1,0}$ | $\psi_{2,0}$ |
|---|---|---|---|---|---|---|
| 12A–86G,(1) | 7.1338 | 2.4933 | 2.5438 | 2.6899 | 0.2572 | -0.1774 |
| 16G–84A,(1) | 7.8676 | 1.6454 | 1.2439 | 0.2123 | 1.3383 | 1.6159 |
| 17G–56A,(1) | 6.869 | 1.2542 | 1.7493 | -2.926 | 1.5819 | 2.0062 |
| 19G–42G,(2) | 6.9976 | 1.4496 | 1.4946 | -2.5631 | 2.0442 | 1.4563 |
| 19G–47A,(2) | 6.6189 | 2.9945 | 2.7491 | 1.2602 | 1.1617 | 0.7671 |
| 28U–32U,(2) | 5.3492 | 2.6351 | 2.9348 | 2.3064 | 0.521 | 1.8542 |
| 37G–69A,(1) | 7.517 | 1.4115 | 2.8021 | 2.2299 | 1.5437 | 1.654 |
| 53A–84A,(2) | 6.1941 | 2.244 | 1.7959 | 2.5803 | 2.615 | -0.9895 |
| 56A–83G,(2) | 6.7146 | 1.9838 | 1.3896 | 2.8881 | -1.4999 | 1.8251 |
| 60G–78G,(2) | 6.6623 | 2.8644 | 2.0505 | -0.0205 | 1.8155 | -2.9846 |
| 61A–77C,(1) | 7.1832 | 2.9863 | 1.7186 | -2.4062 | -0.6842 | 1.662 |

**Table S6: Nucleotide number of base-pair forming $\pi - \pi$ tertiary stack in TPP AD.**

Parameters are derived from coarse-grained structure of TPP AD.

| $B_1$–$B_2$ | $r_0$ | $\theta_{1,0}$ | $\theta_{2,0}$ | $\psi_0$ | $\psi_{1,0}$ | $\psi_{2,0}$ |
|---|---|---|---|---|---|---|
| 24–69 | 3.5188 | 1.8007 | 1.8291 | -0.3482 | -2.5267 | 2.7887 |
